# Decoding Condensate Composition Reveals Tunable Interactomes and Functions of Disordered Proteins

**DOI:** 10.1101/2025.10.27.684971

**Authors:** Qinyu Han, Yanghao Zhong, Baiyi Quan, Ting-Yu Wang, Tsui-Fen Chou, Shasha Chong

**Affiliations:** Division of Chemistry and Chemical Engineering, California Institute of Technology, Pasadena, CA 91125, USA; Proteome Exploration Laboratory, Beckman Institute, California Institute of Technology, Pasadena, CA 91125, USA; Division of Biology and Biological Engineering, California Institute of Technology, Pasadena, CA 91125, USA

**Author notes:** These authors contributed equally (Yanghao Zhong, Baiyi Quan).

## Abstract

Many intrinsically disordered protein regions (IDRs) can form liquid-liquid phase separation (LLPS) condensates via multivalent interactions, but how the interaction behaviors contribute to specific cellular functions remains unclear due to the unknown composition of the condensates. Here, we report **phase**-separation-induced **i**nteractome **d**etection (PhaseID), a new method that determines the protein components within the LLPS condensates driven by any given IDR in live human cells. Using PhaseID, we demonstrated that transactivation IDRs each have unique multivalent interactomes, thereby executing transactivation through distinct pathways. In-depth analyses of PhaseID-determined IDR interactomes revealed that LLPS tunes the interaction selectivity of IDRs, enabling them to play new roles in specific cellular processes, and different LLPS condensates recruit proteins with distinct physicochemical properties encoded in their sequences.

## Main Text

Often referred to as the “dark proteome”, intrinsically disordered protein regions (IDRs) do not fold into well-defined 3D structures and are therefore not amenable to classical structure-function analysis. Although IDRs constitute 44-54% of the eukaryotic and viral proteomes (*1*), play essential roles in numerous cellular processes (*2*), and contain over 20% of all disease mutations (*3*), how most IDRs perform specific functions remains unclear. This void represents a significant hurdle for understanding pathological mechanisms and developing therapeutics for numerous human diseases. The past 15 years have witnessed the groundbreaking discovery that many IDRs can form liquid-liquid phase separation (LLPS) condensates via their multivalent interactions, i.e., protein-protein interactions with a variable, concentration-dependent stoichiometry (*4*–*6*). An explosive number of reports have documented that such interaction behaviors play important roles in cellular organization and many cellular functions (*7*–*13*). These works have opened a new frontier for studying IDRs, but substantial challenges remain in understanding the mechanisms by which they work. The function of a biomolecule is fundamentally determined by which partners it interacts with and how it interacts with them. Although many recent studies have clearly shown that individual IDRs can have highly selective multivalent interaction partners (*5*, *14*–*17*), there is currently no method capable of identifying all the interaction partners of any given IDR in its LLPS condensates in live cells. It thus remains unclear how most IDRs execute specific cellular functions. Due to the methodological gap, it is equally challenging to uncover exactly how the formation of LLPS condensates and related biomolecular assemblies contributes to cellular functions. We note that IDR-mediated assemblies, regardless of LLPS properties, have been referred to by various names in the literature, including “hubs”, “biomolecular condensates”, “droplets”, and “puncta”, with biomolecular condensates used most frequently. In this paper, when referring to the general context, we use “assemblies” as a neutral, inclusive term that encompasses all intracellular compartments formed via IDR-mediated multivalent interactions, including LLPS condensates. We preserve the original terminology used in the cited studies when discussing them.

Eukaryotic transcriptional regulation exemplifies the various cellular processes involving functional IDRs and IDR-mediated assemblies. Many transcriptional regulators contain sizable IDRs. Preeminent examples are intrinsically disordered transactivation domains of DNA-binding transcription factors (TFs). These TF IDRs mediate interactions between TFs and other regulatory proteins (*18*) to enable gene activation; however, it remains unclear how the IDRs successfully execute such transactivation functions without well-defined protein structures. Previously, using live-cell single-molecule imaging approaches, we found that TF IDRs undergo dynamic, multivalent, and selective interactions to drive the formation of local high-concentration TF hubs at target genomic loci, which can develop into LLPS condensates upon IDR overexpression. The multivalent IDR-IDR interactions play a crucial role in transactivation (*5*). Many other reports have also documented the functional roles of TF hubs or condensates in transcription (see a recent review (*19*)). However, without knowing the composition of a TF hub in live cells, it remains elusive which regulatory proteins a TF coordinates with to orchestrate transactivation and how IDR-mediated TF hubs enable the coordination. Moreover, more than 1,600 human TFs that contain distinct IDRs have been discovered to date (*18*). Whether and how these TF IDRs differ in their transactivation mechanisms is completely unexplored.

To fill these gaps, we establish a method that determines the protein components within the LLPS condensates driven by any given IDR in live human cells. This new method, <u>Phase</u>-separation-induced Interactome <u>D</u>etection (PhaseID), combines chemical induction of LLPS with proximity labeling mass spectrometry to enable high-throughput identification of a target IDR’s multivalent protein partners *in situ* within its LLPS condensates in live cells. PhaseID is generally applicable to any IDR with an LLPS propensity and compatible with diverse cell types, providing a useful tool for understanding many cellular processes involving IDRs or biomolecular condensates.

Here, we use PhaseID to study human TF IDRs, validate the PhaseID results with orthogonal assays, and dissect how the IDRs’ selective multivalent interactions contribute to transcriptional regulation. Specifically, we compare the PhaseID-determined multivalent interactomes of different TF IDRs and examine the functional roles of PhaseID-detected IDR partners. In addition, we explore the origin of proteins’ selectivity in interacting with specific IDRs by computing diverse sequence-encoded physicochemical properties of the proteins enriched in and depleted from different LLPS condensates. We have found that the investigated TF IDRs each have signature multivalent interactomes and execute transactivation by interacting with distinct protein partners. An IDR can alter its interaction selectivity and participate in new cellular functions when it undergoes LLPS. We have also shown that LLPS condensates driven by different IDRs recruit proteins with distinct sequence features.

### PhaseID determines an IDR’s multivalent interactome in its LLPS condensates in live human cells

The challenge in characterizing the composition of LLPS condensates stems from the fact that multivalent interactions between IDRs, which drive condensate formation, are weak, transient, and highly sensitive to protein concentrations and other environmental parameters (*2*). These features are distinct from stoichiometric interactions between structured proteins. Thus, LLPS condensates formed in live cells are not readily preserved during biochemical purification and are not amenable to composition characterization *in vitro,* as the environment is drastically different from that within the cell. PhaseID overcomes these challenges by using a novel way to chemically induce LLPS of a target IDR in live cells and determining the protein components within the induced condensates *in situ*.

Specifically, PhaseID utilizes a chemically inducible heterodimerization system comprising FKBP and FRB, where the two proteins bind in the presence of rapalog (a rapamycin analog) (*20*). The FKBP-FRB heterodimer associates an IDR of interest with FTH1, a 24-mer-forming protein (*21*), only in the presence of rapalog, which promotes LLPS of the IDR by increasing its interaction valence. The chemically induced heterodimer, meanwhile, associates the IDR with the biotin protein ligase TurboID, which can biotinylate proteins in its proximity (*22*). As a result, TurboID biotinylates all proteins interacting with the IDR, which is primarily located in its LLPS condensates. Using rapalog (a small molecule) to control LLPS allows the acquisition of a homogeneous cell population, where each cell contains many condensates of the target IDR upon chemical induction, and the system readily scales to large numbers of cells. Obtaining a sizable cell population is crucial for maximizing the detection sensitivity of in-condensate proteins, as many sizable condensates in each cell can significantly enrich these proteins, and many such cells ensure the production of a sufficiently large biotinylated protein sample for mass spectrometry. As such, the workflow of PhaseID begins with stable expression of two proteins, “core” and “bait”, in the cell. The core protein is a fusion of FKBP, TurboID, EGFP, and FTH1, while the bait protein is a fusion of FRB, mCherry, and the target IDR. We next treat the cells with rapalog to induce LLPS of the target IDR, which can be monitored by fluorescence imaging of EGFP and mCherry. We then add biotin to the cells to enable TurboID-mediated biotinylation of all proteins in the induced LLPS condensates, purify the biotinylated proteins using streptavidin-coated beads, and identify the proteins using quantitative mass spectrometry (Fig. 1A).

**Fig. 1.**
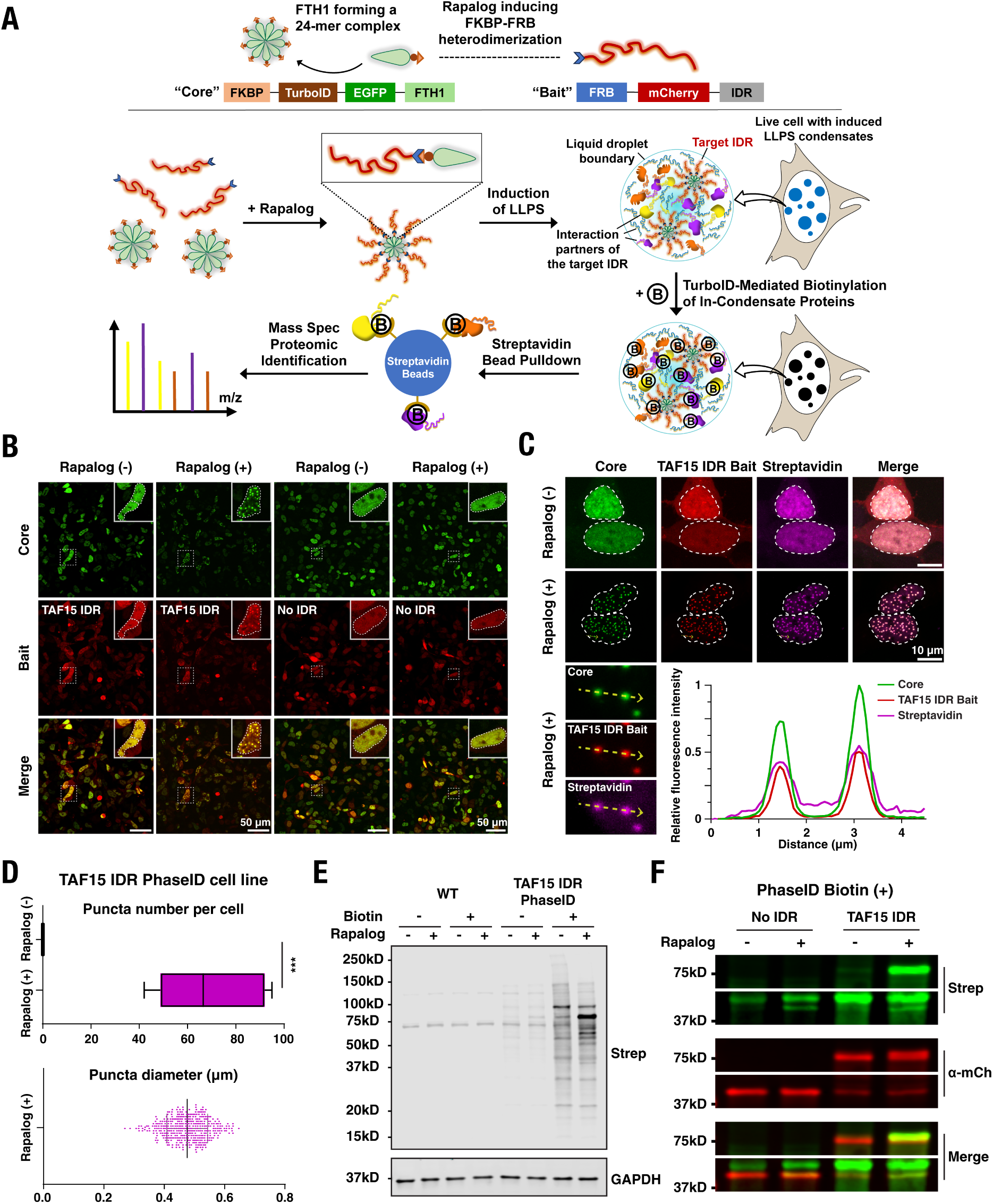
PhaseID enables chemically induced LLPS and specific in-condensate biotinylation in live cells. (A) Schematic of the PhaseID workflow. We stably express two fluorescently tagged fusion proteins, core and bait, in the cell. LLPS of the IDR of interest occurs upon rapalog addition to the cells, which induces the association of core and bait via FKBP-FRB heterodimerization. TurboID mediates proximity biotinylation of proteins within the LLPS condensates. The biotinylated proteins are then isolated by streptavidin pull-down and identified by mass spectrometry. We also perform a control PhaseID without an IDR in the bait to identify the proteins binding to the domains of core and bait other than the target IDR. (B) Confocal fluorescence images of modified HEK293 cells expressing the TAF15 IDR or no-IDR control PhaseID constructs before and after rapalog treatment (500 nM, 10.5 hrs). Rapalog induces LLPS of TAF15 IDR in live cells. The distributions of core and bait are visualized in the EGFP (green) and mCherry (red) channels, respectively. A representative cell under each condition is zoomed in. A white dashed contour outlines the cell nucleus. Scale bars, 50 µm. (C) Higher-magnification views of rapalog-induced TAF15 IDR condensates in HEK293 cells. After the cells were treated with biotin (50 µM, 1 hr), cell staining with AF750-labeled streptavidin (magenta) showed that protein biotinylation predominantly occurs in the condensates. A white dashed contour outlines the cell nucleus. Scale bars, 10 µm. Line profiles show a strong co-localization of the core, bait, and streptavidin signals within the condensates. (D) Quantification of puncta number per cell (top) and puncta diameter (bottom) across six cells in the AF750-labeled streptavidin channel, showing rapalog-dependent TAF15 IDR condensate formation. The number of puncta per cell is presented as a Tukey box plot. Each dot in the bottom plot represents a punctum. Statistical significance: *p* < 0.001 (***); Welch’s two-sample *t*-test. (E) Streptavidin Western blot (Strep) of whole-cell lysates from wild-type and TAF15 IDR PhaseID cells before and after treatment of rapalog (500 nM, 10.5 hrs) and biotin (50 μM, 1 hr). The overall biotinylation levels in TAF15 IDR PhaseID cells increase significantly upon rapalog-induced LLPS. GAPDH served as a loading control. (F) Streptavidin (Strep) and anti-mCherry (α-mCh) blots showing rapalog-dependent biotinylation of the mCherry-labeled TAF15 IDR bait (∼61.7 kDa) and no-IDR bait (∼39.4 kDa), confirming successful core-bait association upon chemical induction of LLPS. Equal amounts of total protein were loaded in all lanes, as verified by Ponceau S staining of the same membrane.

We first demonstrated PhaseID in HEK293 cells using the N-terminal IDR of TAF15 (*23*) (residues 2-205; referred to as TAF15 IDR below), which is known to undergo LLPS under specific conditions (*24*–*26*). TAF15 IDR can execute transactivation when fused to a DNA-binding domain (*23*). Since transactivation occurs in the nucleus, and to enable comparison of the multivalent interactomes of TAF15 IDR and other transactivating IDRs, we performed PhaseID for these IDRs in the nucleus by fusing nuclear localization signal (NLS) peptides to both core and bait (fig. S1A). Whereas both the core and TAF15 IDR-containing bait were distributed homogeneously in the nucleoplasm of live HEK293 cells, the two proteins co-condensed into numerous prominent puncta in the nucleus when we treated the cells with 500 nM rapalog (Fig. 1B). These puncta displayed hallmark features of LLPS condensates, including concentration-dependent formation, spherical morphology, and fusion upon contact. Approximately 70% of the cells contained such nuclear puncta. As a control, we removed TAF15 IDR from the bait and kept all the other settings unchanged (fig. S1A). We observed no puncta formation after treating the cells expressing the “no IDR” bait with rapalog (Fig. 1B). These results suggest that the rapalog-induced puncta in the TAF15 IDR-bait-expressing cells are indeed LLPS condensates driven by TAF15 IDR.

We next incubated the rapalog-treated cells with 50 μM biotin for 1 hour to enable TurboID-mediated proximity labeling. We found that protein biotinylation predominantly occurs in the induced TAF15 IDR condensates, as determined by cell staining with fluorescently labeled streptavidin (Fig. 1C). Each cell has 69 ±22 biotinylated TAF15 IDR condensates with an average diameter of 0.476 ± 0.0679 μm (Fig. 1D). In contrast, cells expressing the control bait with no IDR have biotinylated proteins distributed nearly homogeneously with no detectable puncta throughout the nuclei (fig. S1B). Using Western blots, we observed an increased overall level of protein biotinylation in the TAF15 IDR-bait-expressing cells after rapalog treatment (Fig. 1E). The newly biotinylated proteins included the mCherry-labeled TAF15 IDR bait (Fig. 1F). Similarly, in cells expressing the control bait with no IDR, the bait was biotinylated only after the cells were treated with rapalog (Fig. 1F), though the overall protein biotinylation level only minimally increased compared with the TAF15 IDR-bait-expressing cells upon rapalog treatment (fig. S1D). By driving LLPS, TAF15 IDR recruits many protein molecules into its condensates and enables their biotinylation, thereby increasing overall biotinylation levels in the cell. These results are consistent with our expectation that, only upon rapalog treatment that induces core-bait association, the bait and its interaction partners become proximal to TurboID and get biotinylated. Regardless of rapalog treatment and the presence of TAF15 IDR in the bait, the GFP-labeled core that contains TurboID is always biotinylated as expected (fig. S1F).

To identify proteins whose biotinylation levels are sensitive to LLPS induction by rapalog, we performed proximity labeling and mass spectrometry under both rapalog induction and no rapalog conditions. We then calculated the fold change in the abundance of each detected biotinylated protein upon rapalog induction relative to the no rapalog condition, and identified proteins with a fold change above 1.5 or below 0.67, with statistical significance, as enriched in or depleted from the induced TAF15 IDR condensates, respectively. Since proteins can be enriched in the condensates through interactions with other domains in the core and bait besides TAF15 IDR, we performed a control PhaseID in cells expressing the bait with no IDR to identify the nonspecific partners. Proteins enriched or depleted in this control were removed from the condensate-enriched or depleted hits in the TAF15 IDR PhaseID, respectively, to define the multivalent interactome of TAF15 IDR shown in fig. S2.

### Orthogonal fluorescence imaging assays validate the PhaseID-determined multivalent interactome

We employed cell imaging approaches orthogonal to mass spectrometry to validate the results obtained with PhaseID. Since PhaseID requires overexpressing the target IDR and associating it with several other proteins (mCherry, EGFP, FTH1, and the FKBP-FRB heterodimer), making an environment different from that of the endogenous IDR-containing protein, it is critical to confirm that the PhaseID-detected protein-protein interactions occur between endogenous proteins in the cell. We previously generated reagents to image the interaction behavior of an endogenous IDR-containing protein, EWS::FLI1. It is an oncogenic TF known to cause Ewing sarcoma (*27*) and produced by chromosomal translocation that fuses the N-terminal IDR of EWSR1 (*28*), an RNA-binding protein, to the C-terminal DNA-binding domain of FLI1, a TF that plays roles in development and homeostasis (*29*). We performed PhaseID for EWS IDR (residues 2-264 of EWS::FLI1; referred to as EWS IDR below) and leveraged our previously established cell imaging assays to evaluate the PhaseID-identified partners of EWS IDR under both overexpression and native conditions.

Similar to TAF15 IDR PhaseID, we successfully induced LLPS of EWS IDR in the nucleus of live HEK293 cells by treating the cells with 500 nM rapalog. Approximately 90% of the cells contained nuclear puncta (Fig. 2A). We then acquired the multivalent interactome of EWS IDR (Fig. 2B). We found that PhaseID identified multiple known interaction partners of EWSR1 or EWS::FLI1 (*5*, *30*–*34*) (Fig. 2B). We then employed the “LacO array assay” (*5*), which can characterize the multivalent interactions between any pair of proteins by quantitative cell imaging, to validate new EWS IDR partners identified by PhaseID. The LacO array assay exploits a large synthetic array of lac operators (LacOs) incorporated into the genome of U2OS cells (*35*). The LacO array becomes a local high-concentration region for any IDR of interest when it is fused to lac repressor (LacI) and EYFP and expressed in the cell. To detect potential interactions between the IDR and another protein of interest (POI), we co-express POI labeled with mCherry in the cell and examine whether mCherry-POI is enriched at the LacO array bound by EYFP-IDR-LacI using fluorescence microscopy (Fig. 2C). We also examine potential enrichment of mCherry-POI at the LacO array bound by EYFP-labeled LacI only. A higher mCherry-POI enrichment at the LacO array bound by EYFP-IDR-LacI than by EYFP-LacI would suggest IDR-POI interactions. The degree of mCherry-POI enrichment at the LacO array bound by EYFP-IDR-LacI is correlated with the IDR-POI interaction affinity. Using the LacO array assay, we confirmed the interactions between EWS IDR and several PhaseID-identified EWS IDR partners, including ARID1A (*36*), LDB1 (*37*), LIMD1 (*38*), NONO (*39*), SUMO2 (*40*), and YAP1 (*41*) (Fig. 2D). We observed that these partners interact with EWS IDR with different affinities, as shown by various degrees of mCherry enrichment at the LacO array (Fig. 2E). We also confirmed that ARID4B (*42*) and ZBED4 (*43*), proteins depleted from the induced EWS IDR condensates in PhaseID, do not interact with EWS IDR (Fig. 2D and E). We note that the PhaseID hits examined using the LacO array assay were randomly selected from the lists of enriched and depleted proteins. These results demonstrate the validity of PhaseID in determining the multivalent interactome of a target IDR.

**Fig. 2.**
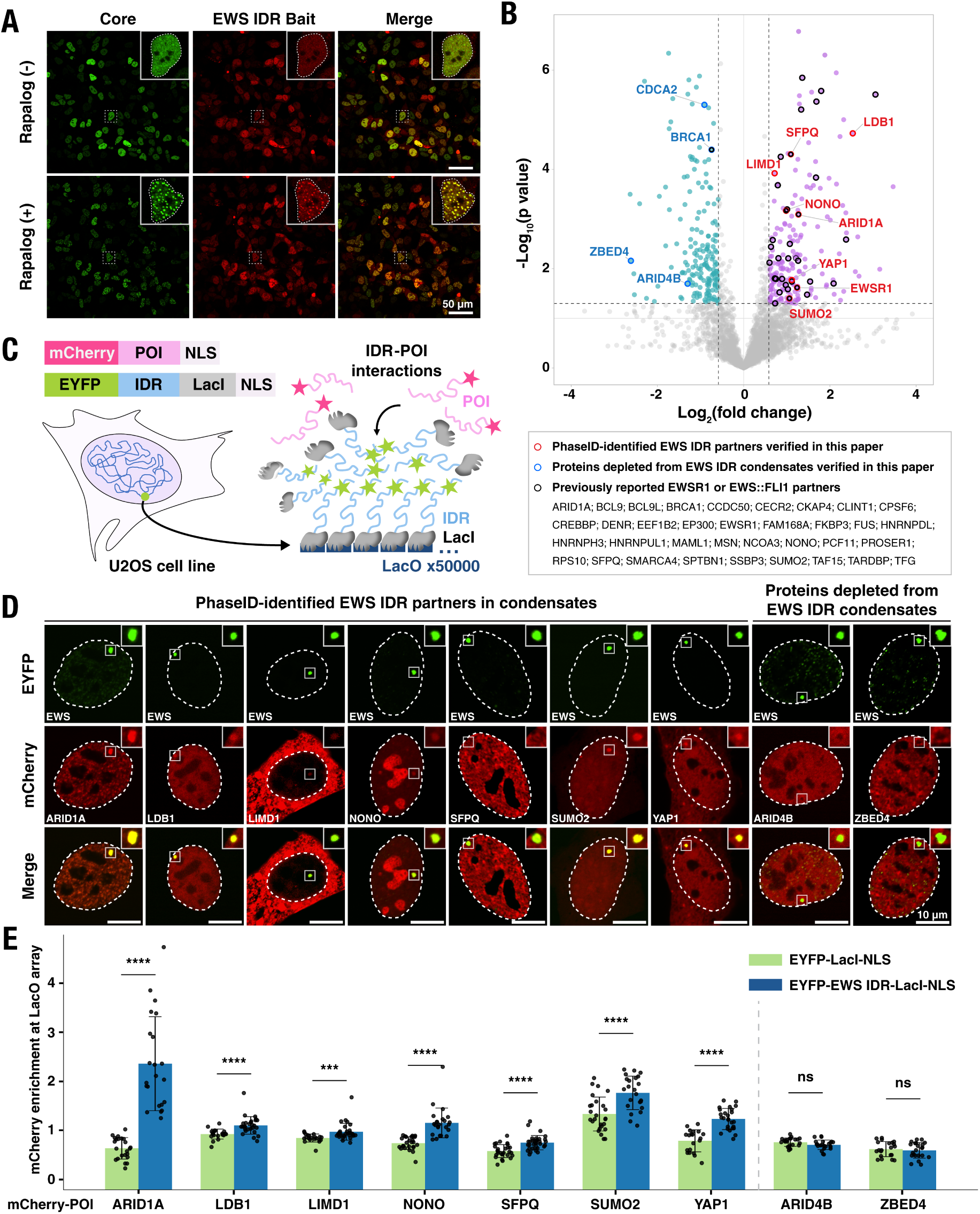
PhaseID determines the interactome of EWS IDR, which is validated by the LacO array assay. (A) Confocal fluorescence images of modified HEK293 cells expressing the EWS IDR PhaseID constructs before and after rapalog treatment (500 nM, 2 hrs). Rapalog induces LLPS of EWS IDR in live cells. The distributions of core and bait are visualized in EGFP (green) and mCherry (red) channels, respectively. A representative cell under each condition is zoomed in. A white dashed contour outlines the cell nucleus. Scale bars, 50 µm. (B) Volcano plot showing the PhaseID-determined EWS IDR interactome, based on nine biological replicates per condition (before and after rapalog treatment). Each protein is represented by a dot in the plot. Proteins significantly enriched in and depleted from the rapalog-induced EWS IDR condensates are shown in magenta and blue, respectively. Proteins verified in this study using orthogonal assays are highlighted with red or blue circles. Previously reported EWSR1 or EWS::FLI1 partners are highlighted with black circles (*117*, *118*). The x-axis represents the fold change of the detected protein abundance upon rapalog induction compared with no rapalog condition. The y-axis represents the significance of the fold change. The dashed horizontal line shows the threshold *p* value of statistical significance (0.05). Statistical significance was determined by an unpaired Student’s *t*-test. The two dashed vertical lines show the thresholds of abundance fold change (1.5 or 0.67). (C) Schematic of the LacO array assay where we express EYFP-IDR-LacI and mCherry-tagged protein of interest (POI) in the LacO-containing U2OS cells. We fuse an NLS peptide to each of the proteins to ensure their nuclear localization. Colocalization of the two proteins at the LacO array suggests LCD-POI interactions. (D) Confocal fluorescence images of LacO-containing U2OS cells co-expressing EYFP-EWS IDR-LacI-NLS and mCherry-labeled proteins identified by PhaseID. The region surrounding the LacO array is zoomed in. Scale bars, 10 µm. (E) Quantification of mCherry-POI-NLS enrichment at the LacO array bound by EYFP-LacI-NLS or EYFP-EWS IDR-LacI-NLS. The enrichment was calculated as the peak mCherry fluorescence at the array divided by the average intensity immediately surrounding the array. Higher enrichment of mCherry-POI-NLS at the array bound by EYFP-EWS IDR-LacI-NLS than by EYFP-LacI-NLS suggests EWS IDR–POI interactions. Error bars represent standard deviations. Statistical significance: *p* < 0.001 (***), *p* < 0.0001 (****), not significant (ns); Welch’s two-sample *t*-test.

We next examined whether PhaseID-identified EWS IDR partners interact with endogenous EWS::FLI1 in the Ewing sarcoma cell line A673. We utilized a knock-in A673 cell line that we previously generated using CRISPR-mediated genome editing, in which a HaloTag labels endogenous EWS::FLI1 without disrupting its functions (*5*). Since HaloTag can covalently bind to a fluorescent ligand with excellent photophysical properties (*44*, *45*), such labeling allows live-cell imaging of endogenous EWS::FLI1 at its native expression levels and with single-molecule resolution. We previously visualized the formation of small, transient, and EWS IDR-mediated EWS::FLI1 hubs at endogenous target genes in these cells (*5*). Here, we focused on several PhaseID-identified EWS IDR interaction partners (YAP1, EWSR1, SFPQ (*46*), and SUMO2) and imaged each endogenous protein by immunofluorescence while simultaneously imaging endogenous EWS::FLI1-Halo (Fig. 3A and fig. S4A). We detected significant enrichment of each partner at endogenous EWS::FLI1 hubs (Fig. 3B and C). In contrast, BRCA1 (*47*) and CDCA2 (*48*), proteins depleted from the induced EWS IDR condensates in PhaseID, were not enriched in endogenous EWS::FLI1 hubs (Fig. 3B and C, and fig. S4A). We note that our choice of the examined PhaseID hits was based on our access to reliable, immunofluorescence-compatible antibodies that target these proteins. Our results suggest that PhaseID, despite involving target IDR overexpression in the workflow, detects protein-protein interactions of the endogenously expressed target IDR in its native cellular context. As such, PhaseID determines functionally relevant multivalent interactomes.

**Fig. 3.**
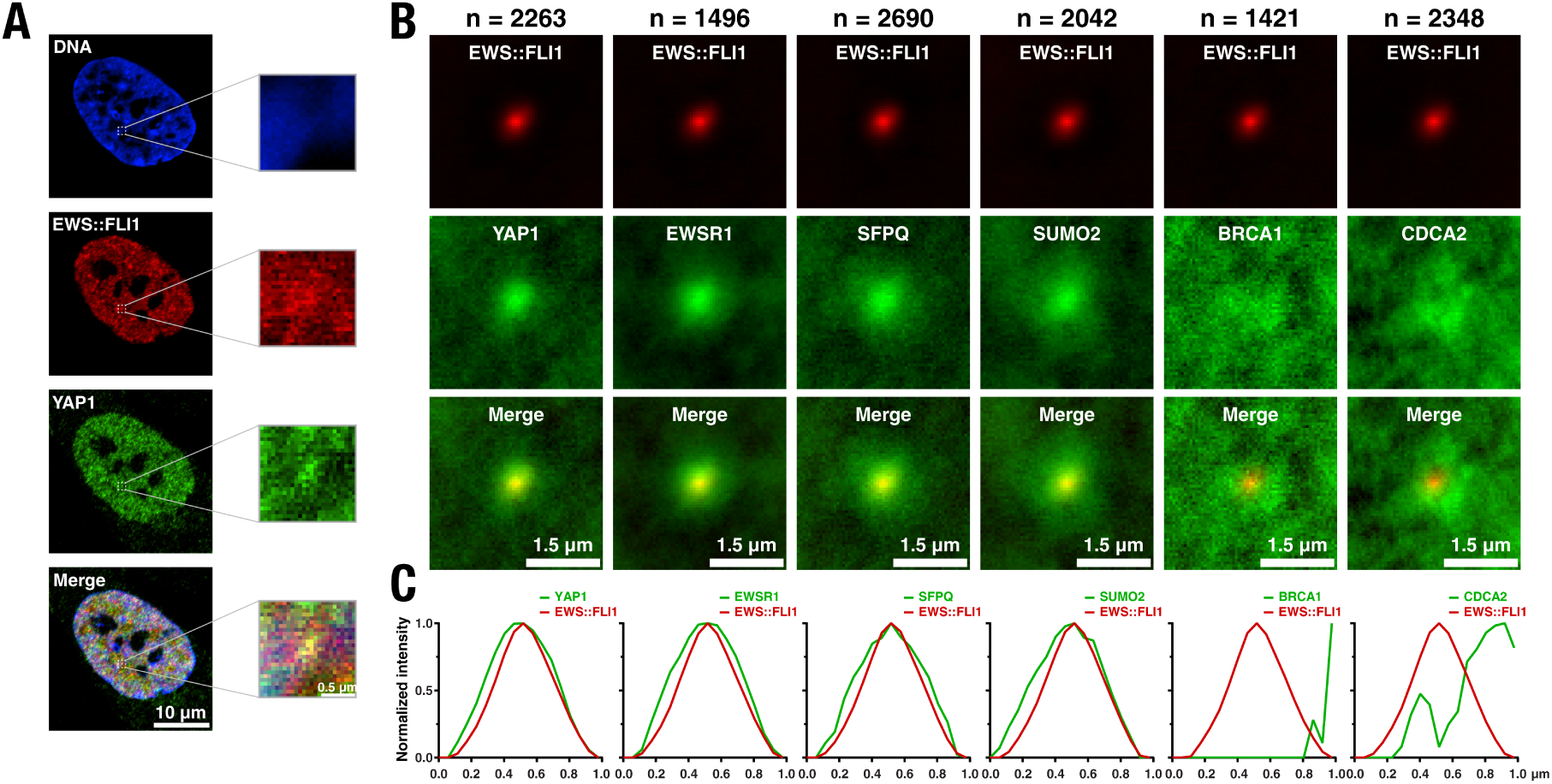
PhaseID-identified EWS IDR interaction partners are enriched in endogenous EWS::FLI1 condensates. (A) Confocal fluorescence images of a knock-in A673 cell nucleus showing DNA stained with Hoechst (blue), endogenous EWS::FLI1-Halo labeled with the HaloTag ligand JFX646 (*119*) (red), and endogenous YAP1 immunostained with a primary YAP1 antibody and an AF488-labeled secondary antibody (green). The zoom-in images show a single EWS::FLI1 hub and fluorescence of the same field of view in all channels. (B) Averaged two-color confocal fluorescence images showing endogenous EWS::FLI1-Halo hubs (red) and immunostained endogenous proteins (green) detected by EWS IDR PhaseID. Four EWS IDR partners (YAP1, EWSR1, SFPQ, and SUMO2) and two proteins depleted from the EWS IDR condensates (BRCA1 and CDCA2) are investigated. The number of analyzed EWS::FLI1 hubs for each protein is provided above the respective two-color image (11–20 cells analyzed for each protein). (C) Normalized intensity profiles of the central region (1 µm × 1 µm square) of the two-color fluorescence images in (B). The profiles were generated by averaging pixel intensities along each column within the central region.

### Multivalent interactomes of transactivating IDRs indicate different transactivation mechanisms

We applied PhaseID to understand how different IDRs execute transactivation. To this end, we measured the multivalent interactomes of three transactivating IDRs, i.e., TAF15 IDR, EWS IDR, and the IDR of Sp1 (residues 2-507; referred to as Sp1 IDR below), a TF involved in regulating many cellular processes (*49*, *50*). Similar to TAF15 and EWS IDRs, we induced LLPS of Sp1 IDR in the nucleus of live HEK293 cells with 500 nM rapalog (fig. S5B and C) and determined its multivalent interactome in the induced condensates (fig. S5D). By comparing the PhaseID results for all three IDRs, we found that their interactomes are distinct from each other with only small overlaps (Fig. 4A). 284 out of the 348 interaction partners of TAF15 IDR are unique to TAF15 IDR, 141 out of the 204 partners of EWS IDR are unique to EWS IDR, and 92 out of the 120 partners of Sp1 IDR are unique to Sp1 IDR. The IDR interactomes overlap to different degrees. EWS IDR shares 50 partners with TAF15 IDR and only 14 partners with Sp1 IDR. TAF15 IDR only shares 15 partners with Sp1 IDR. This result suggests that Sp1 IDR has a multivalent interactome that is most distinct among the three, whereas the interactomes of EWS and TAF15 IDRs are more similar to each other despite their differences. We also found that endogenous FET family proteins (TAF15, FUS, and EWSR1), but not Sp1, are significantly enriched in the EWS and TAF15 IDR condensates. In contrast, endogenous Sp1 but not TAF15, FUS, or EWSR1 is significantly enriched in the Sp1 IDR condensates (fig. S6). This result is consistent with previously reported findings by us and others, which show that Sp1 homotypically interacts, and the FET family proteins homotypically and heterotypically interact within the family, all via their IDRs, but Sp1 does not interact with the FET family proteins (*5*, *26*). The successful detection of the expected IDR-mediated interactions provides additional validation of the PhaseID-determined interactomes.

**Fig. 4.**
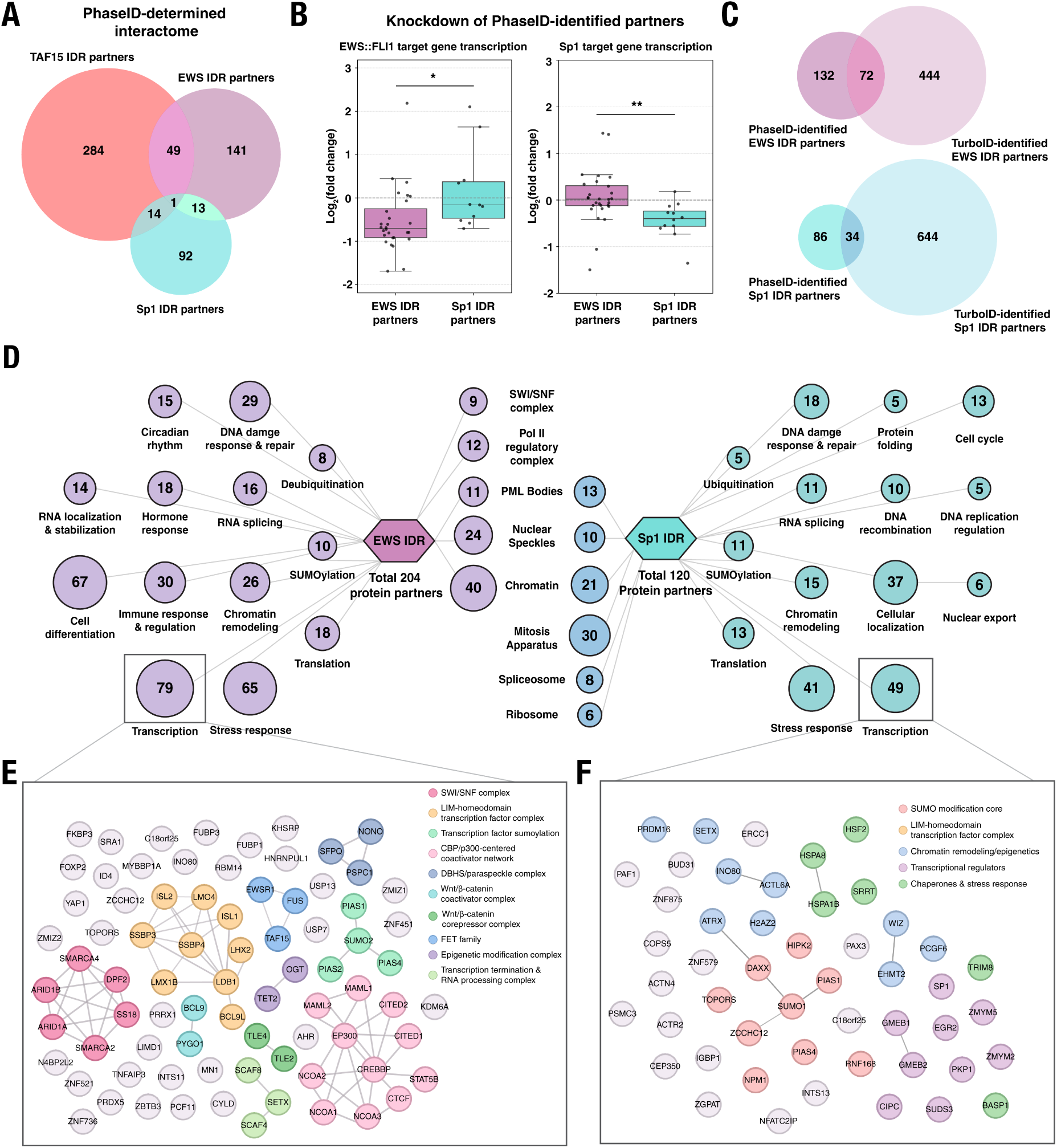
PhaseID reveals that different transactivation IDRs have distinct, functionally relevant multivalent interactomes. (A) Venn diagram comparing the PhaseID-determined interactomes of TAF15, EWS, and Sp1 IDRs. (B) siRNA-mediated knockdown of PhaseID-identified IDR partners in A673 cells specifically affects transcription activated by the corresponding IDR-containing transcription factor. Left, knocking down EWS IDR partners (SFPQ, TAF15, SUMO2, LDB1, HNRNPUL1, MN1, or LIMD1) but not Sp1 IDR partners (ZMYM2, HSF2, or PCGF6) reduces transcription of EWS::FLI1 target genes (*CYP4F22*, *CCK*, *PPP1R1A*, and *NR0B1*). Right, knocking down Sp1 IDR partners but not EWS IDR partners listed above reduces transcription of Sp1 target genes (*BIRC5*, *B4GALT5*, *MMP14*, and *IGF1R*). The transcription levels were measured by RT-qPCR. The y-axis represents the fold change in transcription level upon siRNA knockdown compared with that in A673 cells transfected with control siRNA, which has no specific target in the cell. Data presented as Tukey box plots. Each dot represents the fold change of each target gene upon knockdown of one partner. Statistical significance: *p* < 0.05 (*)*, p* < 0.01 (**); Welch’s two-sample *t*-test. (C) Venn diagrams comparing the PhaseID- and TurboID-determined interactomes of EWS IDR (top) and Sp1 IDR (bottom). (D) Gene Ontology (GO) enrichment analysis of the PhaseID-identified EWS IDR (left) and Sp1 IDR (right) interaction partners. The center nodes represent significantly enriched “Cellular Component” terms among the IDR partners, and the side nodes represent enriched “Biological Process” terms among the IDR partners. (E-F) Network representation of PhaseID-identified EWS IDR partners (E) and Sp1 IDR partners (F) that are associated with transcription in the GO enrichment analysis.

The fact that the three transactivating IDRs have distinct multivalent interactomes supports a model in which the IDRs each enable the corresponding TFs to form a signature “flavor” of hubs with a unique composition, recruiting a particular group of regulatory proteins to target genes and executing transactivation through a distinct pathway. Based on this new model, we hypothesize that a protein uniquely interacting with the IDR of one TF only contributes to the functions of that TF and does not affect the functions of other TFs with which it does not interact. We focused on two TFs, EWS::FLI1 and Sp1, to test this hypothesis. Specifically, we examined how the gene activation functions of EWS::FLI1 and Sp1 change in A673 cells, where both TFs are expressed endogenously, upon knockdown of specific PhaseID-identified interaction partners of EWS IDR or Sp1 IDR. The investigated protein partners unique to EWS IDR were SFPQ, TAF15, SUMO2, LDB1, HNRNPUL1 (*51*), MN1 (*52*), and LIMD1, while those unique to Sp1 IDR were ZMYM2 (*53*), HSF2 (*54*), and PCGF6 (*55*). To characterize the transactivation functions of EWS::FLI1 and Sp1, we focused on their known target genes, i.e., *CYP4F22*, *UGT3A2*, *PPP1R1A*, and *NR0B1* for EWS::FLI1 (*5*, *56*), and *BIRC5*, *B4GALT5*, *MMP14*, and *IGF1R* for Sp1 (*57*–*60*). Using RT-qPCR, we examined how transcription of these genes changes upon siRNA knockdown of each of the above interaction partners of EWS IDR or Sp1 IDR (fig. S7 and S8). We found that transcription of the EWS::FLI1 target genes was significantly reduced upon knockdown of the EWS IDR partners, and it was largely unaffected upon knockdown of the Sp1 IDR partners. On the other hand, transcription of the Sp1 target genes was significantly reduced upon knockdown of the Sp1 IDR partners, and it was unaffected upon knockdown of the EWS IDR partners (Fig. 4B). These results suggest that many PhaseID-identified interaction partners of one TF IDR indeed specifically contribute to the transactivation function of that TF. This finding verifies our hypothesis and supports our model that different TFs can form hubs with distinct compositions, and each TF interacts with a unique group of partners within its hubs to activate genes, as shown in Fig. 6.

### LLPS can tune an IDR’s selectivity in interacting with proteins involved in diverse cellular processes

Previous reports have characterized the interactomes of intrinsically disordered proteins in the cell using classical proximity labeling mass spectrometry (*61*), without localizing the target protein predominantly in its condensates. We asked whether the interactome of an IDR can alter when it undergoes LLPS by comparing IDR interactomes measured by PhaseID and conventional TurboID-mediated proximity labeling mass spectrometry (*62*). For the latter, we fused TurboID and mCherry directly to a target IDR (fig. S9A), stably expressed the fusion protein in HEK293 cells, and performed TurboID-mediated proximity labeling followed by mass spectrometry to biotinylate and identify all the proteins proximal to the IDR. We also performed a “no IDR” control experiment in the cells stably expressing TurboID fused to mCherry only. We calculated the fold change in the abundance of each biotinylated protein in the presence of the target IDR relative to the no IDR control condition. Proteins with significantly higher abundance in the presence of the target IDR were defined as interaction partners of the IDR.

We confirmed that none of EWS, Sp1, and TAF15 IDRs form prominent puncta in the cells expressing the IDR TurboID constructs (fig. S9B). After we added biotin to the cells, TurboID-mediated biotinylation was nearly homogeneously distributed across the cell nucleus (fig. S9C), distinct from the predominantly in-condensate biotinylation distribution in PhaseID (Fig. 1C). We compared the TurboID- and PhaseID-identified interactomes for EWS, Sp1, and TAF15 IDRs. Interestingly, for each IDR, its interactomes identified by TurboID and PhaseID only partially overlap, and a significant portion of its interaction partners identified by one method are not recognized by the other (Fig. 4C). In addition, a subset of protein partners detected by TurboID is excluded from the induced condensates of the target IDR according to PhaseID (fig. S10). This indicates that LLPS of an IDR can tune its interaction selectivity.

To identify all the cellular functions that may be mediated or impacted by LLPS of EWS and Sp1 IDRs, we performed Gene Ontology (GO) enrichment analysis for all PhaseID-identified interaction partners of each IDR using DAVID Bioinformatics Resources (*63*) (Fig. 4D). Besides transcription that is expected to involve many partners of EWS and Sp1 IDRs due to their transactivation functions (Fig. 4E and F), several other biological processes are also overrepresented for the partners of both IDRs, including SUMOylation, chromatin remodeling, stress response, DNA damage response and repair, RNA splicing, and translation. On the other hand, processes such as DNA replication regulation, protein folding, nuclear export, and cell cycle are overrepresented only for Sp1 IDR partners. In contrast, processes such as RNA localization and stabilization, cell differentiation, immune response and regulation, circadian rhythm, and deubiquitination are overrepresented only for EWS IDR partners (Fig. 4D). At the same time, chromatin, nuclear speckles, and PML bodies are overrepresented cellular components for the partners of both EWS and Sp1 IDRs. However, components such as the SWI/SNF complex are overrepresented only for EWS IDR partners, whereas components like the mitotic apparatus and ribosome are overrepresented only for Sp1 IDR partners. These results suggest significant functional differences between EWS and Sp1 IDR condensates. Although Sp1, EWSR1, or EWS::FLI1 have been previously reported to play a role in some of the overrepresented biological processes and cellular components (*64*–*73*), this is the first time that their IDRs are found to interact with specific proteins involved in many of the processes or components in the context of LLPS. While the PhaseID-determined interactions between each IDR and many of its partners still require rigorous validation using orthogonal assays, which is out of the scope of this paper, our work opens the door to future mechanistic studies to understand how Sp1, EWSR1, or EWS::FLI1 may contribute to diverse cellular functions.

To further identify the cellular functions that are particularly sensitive to LLPS, we compared the overrepresented GO terms for the interaction partners of each IDR identified by PhaseID and TurboID. While biological processes including transcription, chromatin remodeling, stress response, DNA damage response and repair, cell cycle, cell differentiation, and RNA splicing stand out for the partners identified by both methods, some processes, such as SUMOylation, translation, and deubiquitination, are overrepresented for the EWS or Sp1 IDR partners identified only by PhaseID (Fig. 4D and fig. S11). This result suggests that the EWS or Sp1 IDR partners involved in these processes interact with the respective IDR only when it forms LLPS condensates or other related assemblies. This highlights the particular importance of LLPS or assembly formation, as the assembly behaviors likely enable new roles for Sp1, EWSR1, or EWS::FLI1 in these processes via IDR-mediated interactions.

For example, deubiquitination is overrepresented among PhaseID-identified EWS IDR partners but not among its TurboID-identified partners. The implication of deubiquitination is consistent with previous works that reported the role of several deubiquitinating enzymes in regulating the stability and oncogenic functions of EWS::FLI1 (*74*–*76*). Here, we detected additional deubiquitinating enzymes in the induced EWS IDR condensates, including USP24 (*77*), USP13 (*78*), UCHL3 (*79*), OTUD5 (*80*), TNFAIP3 (*81*), STAMBPL1 (*82*), and CYLD (*83*). The fact that these enzymes are identified as EWS IDR partners only by PhaseID suggests that they likely interact with EWS::FLI1 only within its hubs (*5*). Future functional studies may reveal a role of these enzymes in regulating EWS::FLI1 specifically in the hubs.

On the other hand, SUMOylation is overrepresented among PhaseID-identified EWS IDR partners and Sp1 IDR partners but not among their respective TurboID-identified partners, suggesting that the involvement of SUMOylation in regulating both IDRs likely requires their LLPS or assembly formation. This finding supports the previously reported functional relationship between protein SUMOylation and LLPS, where SUMOylation within IDRs is implicated in the regulation of LLPS, and LLPS can also facilitate protein SUMOylation (*84*–*86*). While SUMOylation is known to regulate the stability and functions of Sp1 (*87*), it has not been reported to regulate EWSR1 or EWS::FLI1. However, in the induced EWS IDR condensates, we detected multiple proteins involved in SUMOylation, including SUMO2 (*88*), SUMO E3 ligases ZNF451 (*89*) and PIAS1/2/4 (*90*), SIMC1, which contains SUMO-interacting motifs (*91*), and SETX, whose function is critically dependent on its SUMOylation (*92*). Since these proteins are identified as EWS IDR partners only by PhaseID, they likely interact with EWS::FLI1 only within its hubs. Future functional studies may uncover their role in regulating EWS::FLI1 functions in the hubs. Notably, PhaseID identified SIMC1 as the only shared partner of all EWS, Sp1, and TAF15 IDRs; however, TurboID did not detect its interaction with any of the IDRs. It will be of future interest to explore whether SIMC1 plays a role in regulating LLPS in general.

### Proteins recruited to Sp1 IDR and EWS IDR condensates have distinct sequence features

An increasing number of reports have shown that a “sequence grammar” may underlie the interaction selectivity of some IDRs, that is, specific physicochemical properties encoded in the amino acid sequences of an IDR or its protein partners may be required for their selective interactions (*5*, *16*, *17*, *93*). To understand the physicochemical origin of IDR interaction selectivity observed with PhaseID, we computed 140 sequence parameters (*94*–*100*) for each protein that is significantly enriched in or depleted from the induced EWS or Sp1 IDR condensates, compared the parameter values of each condensate-enriched or depleted protein group to those of the mass spectrometry detected proteome, and identified the sequence parameters whose values of the protein groups differed significantly from the proteome values (see Methods for details).

The interesting sequence parameters we identified include the fractions of specific residues and residue patches (consecutive occurrence) as well as parameters describing the charge content and distribution (Fig. 5). First, all condensate-enriched and depleted protein groups have higher fractions of disorder-promoting residues than the proteome, indicating that IDRs of these proteins play a role in their abilities to partition in LLPS condensates selectively. We then examined the sequence parameters of specific protein groups. Compared to the proteome, proteins enriched in Sp1 IDR condensates have higher fractions of charged residues, including aspartic acid, glutamate, and lysine, higher fractions and lengths of DE and DEKR patches (D, aspartic acid; E, glutamate; K, lysine; R, arginine), and an overall higher fraction of charged residues (FCR). These proteins also exhibit higher δ and κ values, suggesting their more uneven charge distributions and greater charge segregation levels than the proteome (*95*). In contrast, the values of the same charge parameters in proteins depleted from Sp1 IDR condensates are lower than or comparable to those in the proteome. Intriguingly, proteins enriched in and depleted from EWS IDR condensates exhibit lower and higher values of these charge parameters, respectively, than the proteome values, showing a trend opposite to that observed for the Sp1 IDR interactome. On the other hand, compared to the proteome, proteins enriched in EWS IDR condensates have higher fractions of specific non-charged residues, including methionine, asparagine, glutamine, glycine, and tyrosine, as well as higher fractions and lengths of Q and NQ patches (Q, glutamine; N, asparagine). Proteins depleted from EWS IDR condensates and enriched in Sp1 IDR condensates both exhibit lower values or no significant difference in these sequence parameters compared to the proteome (Fig. 5). Collectively, these results suggest that proteins selectively recruited to Sp1 IDR and EWS IDR condensates exhibit many opposite sequence features. For each IDR, proteins enriched in and depleted from its condensates often exhibit opposite features as well. In short, the selectivity of proteins in partitioning into specific condensates is strongly correlated with their particular physicochemical properties encoded in the amino acid sequences.

**Fig. 5.**
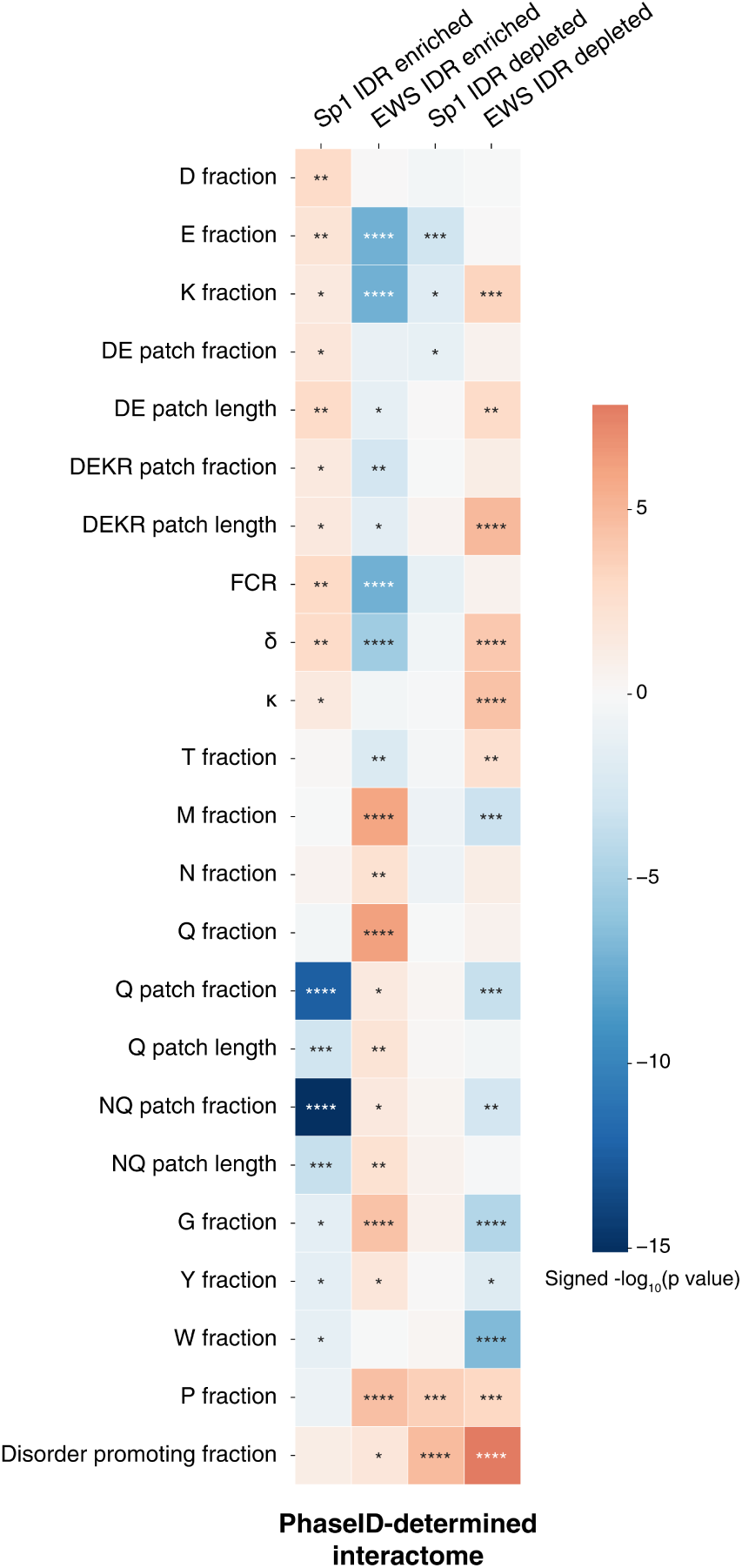
Sequence feature analysis of the PhaseID-determined multivalent interactomes of EWS and Sp1 IDRs. Heatmap of signed *p* values showing that proteins enriched in or depleted from EWS IDR or Sp1 IDR condensates have specific sequence parameter values distinct from those of the mass spectrometry-detected proteome. For each sequence parameter, values for individual PhaseID-identified proteins were normalized to z-scores using the proteome mean and standard deviation. Each z-score represents how far a protein’s parameter value deviates from the proteome average. In the plot, a positive value (red) or a negative value (blue) denotes that the sequence parameter has a higher or lower mean z-score for the protein group, respectively, compared to the proteome. The sign reflects the direction of the mean z-score difference, and the magnitude corresponds to −log₁₀(*p*-value). The sequence parameter values were computed using localCIDER (*94*). FCR, fraction of charged residues. κ quantifies the linear patterning of charged residues along a protein sequence. δ is the variance of local charge asymmetry from the overall charge asymmetry across defined residue windows in a protein sequence. Patches are contiguous runs of ≥5 identical or chemically similar residues. See Methods for details. Statistical significance: *p* < 0.05 (*), *p* < 0.01 (**), *p* < 0.001 (***), *p* < 0.0001 (****); Welch’s two-sample *t*-test.

We next asked whether this sequence feature-dependent interaction selectivity in the PhaseID-determined IDR interactomes is also observed in the TurboID-determined IDR interactomes, where the protein partners do not interact with the target IDR specifically within its LLPS condensates. To this end, we computed 140 sequence parameters for each protein detected by TurboID as interacting with EWS or Sp1 IDR and compared the parameter values of each IDR partner group with those of the mass spectrometry-detected proteome. Although these protein groups show significant deviations from the proteome for many sequence parameters, their sequence features differ markedly from those of PhaseID-identified IDR partners (Fig. 5 and fig. S12). For example, the interaction partners of Sp1 IDR identified by TurboID have lower fractions of lysine and similar fractions of aspartic acid and glutamate compared to the proteome, features distinct from those of proteins enriched in Sp1 IDR condensates identified by PhaseID. Importantly, TurboID-identified partners of Sp1 IDR do not exhibit opposite sequence features to those of EWS IDR partners, as we observed for Sp1 and EWS IDR partners in respective condensates using PhaseID. These findings suggest that the weak, IDR-mediated multivalent interactions occurring specifically in LLPS condensates have the interaction selectivity encoded by a better-defined protein sequence grammar than the IDR-mediated interactions in the absence of condensates. Since the in-condensate interactions can be comprehensively determined by PhaseID but not by conventional proximity labeling mass spectrometry methods, such as TurboID, our findings highlight the unique power of PhaseID in decoding the sequence grammar underlying the selectivity in IDR-mediated multivalent interactions in the context of LLPS or related biomolecular assemblies.

## Discussion

An increasing number of IDRs have been reported to form biomolecular assemblies, including LLPS condensates, via their selective multivalent interactions. Although such interaction behaviors have been clearly shown to be important for the functions of IDRs in many cellular processes, an in-depth mechanistic understanding of how assembly formation contributes to IDR functions is often challenging due to a lack of tools to identify the selective interaction partners of IDRs in the assemblies. Such tools are also highly desirable due to the important roles that LLPS plays in cellular organization and many cellular functions. Identifying the biomolecules assembled into LLPS condensates in the cell is critical for uncovering how exactly LLPS contributes to specific functions. Here, we fill the gaps by establishing the first method (PhaseID) to determine the protein components of LLPS condensates driven by any given IDR in diverse mammalian cell types.

Historically, the challenge in characterizing the composition of LLPS condensates stems from the fact that multivalent interactions between IDRs within condensates, unlike stoichiometric interactions between structured proteins, are weak, transient, and sensitively dependent on protein concentrations and other environmental parameters (*2*). Thus, LLPS condensates formed in live cells cannot be readily preserved during biochemical purification or characterized *in vitro* for their composition, as the environment is drastically different from that within the cell. PhaseID overcomes this obstacle by combining chemical induction of LLPS with proximity labeling mass spectrometry, which enables identifying all the protein components of LLPS condensates *in situ* in live cells. Using a small molecule to induce LLPS allows the acquisition of a large, homogenous cell population where each cell contains many condensates, sufficiently enriching each protein component for mass spectrometry identification.

The experimental design of PhaseID enables advantages over several recent studies that have characterized the composition of specific condensates nucleated by alternative approaches, including *in vitro* droplet formation, optogenetics, and an array of lac operators (LacO). The protocols based on *in vitro* and optogenetic LLPS assays often require significant modification or reinvention for studying each new LLPS-driving protein and are, therefore, not generally applicable methods. For example, determining the composition of MED1 IDR condensates based on *in vitro* droplet formation relies on an essential property of purified MED1 IDR, i.e., it forms LLPS condensates when added to a soluble nuclear extract (*17*). However, many IDRs are difficult to purify, and purified IDRs often require drastically different conditions to undergo LLPS *in vitro* (*101*). As such, many IDRs do not spontaneously form LLPS condensates in the cell extract. Although optogenetic LLPS activation applies to many more IDR-containing proteins, and it, combined with proximity labeling mass spectrometry, has enabled compositional characterization of TopBP1 and RNA Polymerase II condensates (*102*, *103*), light-induced condensates are known to dissolve progressively during the steps of in-condensate protein identification, which can affect the specificity of in-condensate detection, and the protocol of maintaining condensates is dependent on the LLPS-driving protein and needs to be developed on a case-by-case basis (*104*). A modified U2OS cell line containing a LacO array in its genome has been used to nucleate condensates of an IDR of interest when it is fused to lac repressor. This system has recently been combined with proximity labeling mass spectrometry to identify interaction partners of MED1 and FUS IDRs (*105*). Although the LacO array can create a local high-concentration region for any IDR, and thus in principle should allow compositional characterization for nuclear condensates driven by any IDR, the characterization can only be performed in U2OS cells, unless significant efforts are spent in re-establishing a LacO array in the genome of a different cell line. Additionally, there is only one condensate per nucleus nucleated by the LacO array, which occupies a tiny fraction of the nuclear volume and recruits a small number of interaction partners of the target IDR, thereby limiting the downstream detection efficiency of the protein partners by mass spectrometry. Besides, the LacO-array-based method cannot characterize condensates in the cytoplasm.

Distinct from existing methods, PhaseID is designed to be sufficiently robust and sensitive and does not require significant protocol changes to characterize the composition of LLPS condensates driven by a new target protein or in a new cell type. Leveraging chemical induction of many LLPS condensates per cell, PhaseID should detect the target IDR’s protein partners in the condensates with a significantly higher efficiency than the LacO-array-based method. Although in this work, we intentionally induced the formation of nuclear condensates of EWS, TAF15, and Sp1 IDRs because they execute transactivation functions in the nucleus, PhaseID can readily induce LLPS of a target IDR in the cytoplasm and determine the cytoplasmic condensate composition by removing the NLS from core and bait. Overall, the unprecedented versatility and robustness that PhaseID offers will make it universally helpful in studying IDR-mediated biomolecular assemblies across diverse cell types and in understanding how numerous IDR-containing proteins function.

We recognize a limitation of PhaseID - it requires protein overexpression, chemical induction of LLPS, and significantly higher levels of multivalent interactions than those of endogenous proteins. This design ensures high efficiency in recruiting protein partners of the target IDR to its LLPS condensates, thereby maximizing the detection sensitivity of even the weakest interaction partners using downstream proximity labeling mass spectrometry. Due to the synthetic promotion of LLPS in PhaseID, the prominent puncta of a target IDR (Fig. 1B, 2A, and fig. S5B) should not be considered functionally comparable to biomolecular assemblies formed by endogenous proteins in the cell. Our orthogonal cell imaging experiments have demonstrated that PhaseID-detected interactions occur between endogenous proteins (Fig. 3). Novel microscopy approaches, such as proximity-assisted photoactivation (PAPA), which is capable of measuring IDR-mediated interactions between endogenous proteins with single-molecule resolution in live cells (*106*), may provide a valuable tool to further validate PhaseID-identified IDR partners in the future.

One of the IDRs we studied using PhaseID is EWS IDR. It is part of the oncogenic TF EWS::FLI1 that drives the development of Ewing sarcoma, an aggressive and metastatic pediatric cancer that is poorly managed with standard treatments. The survival rate for Ewing sarcoma patients with metastases or recurrence remains extremely low (*27*). While several targeted therapies for Ewing sarcoma have been proposed over the past two decades (*107*), none of them have shown promising results in clinical trials. New, effective therapeutic strategies are thus urgently needed. EWS::FLI1 is long considered an attractive yet notoriously difficult target for drug development as it plays a critical role in driving the oncogenic transcription program in Ewing sarcoma (*108*), though with mechanisms insufficiently understood, mainly due to its disordered nature. We previously showed that EWS::FLI1 forms pathological hubs at its target genes via multivalent interactions of EWS IDR, and this interaction behavior plays a crucial role in both transcription and transformation functions of EWS::FLI1 (*5*, *109*). However, the compositional characterization of the EWS::FLI1 hubs was lacking due to a lack of suitable tools. Here, we fill in the gap by using PhaseID to identify the multivalent interaction partners of EWS IDR in its induced LLPS condensates in HEK293 cells and validating the presence of multiple identified partners in endogenous EWS::FLI1 hubs in Ewing sarcoma cells. In the future, it will be of great interest to comprehensively validate all the PhaseID-detected EWS IDR partners in terms of their interactions with endogenous EWS::FLI1 and the roles they play in EWS::FLI1 functions in Ewing sarcoma cells. Together, the current and future works hold the promise of identifying new essential regulatory factors of EWS::FLI1 and new therapeutic targets for Ewing sarcoma that are otherwise inaccessible, which will pave the way for the development of novel therapeutics. Moreover, aberrant IDR-containing proteins and IDR-mediated pathological assemblies are involved in numerous diseases besides Ewing sarcoma (*110*–*113*). Using PhaseID to characterize the composition of these pathological assemblies in the future will likely shed new light on both the mechanisms and therapeutics of these diseases.

Leveraging the versatility of PhaseID, we found that three transactivating IDRs have distinct multivalent interactomes within their respective LLPS condensates. In particular, focusing on EWS and Sp1 IDRs, we found that many PhaseID-identified, unique partners of each IDR specifically contribute to the transactivation function of the corresponding TF. These results suggest that TFs may form IDR-mediated hubs with distinct compositions, thereby executing transactivation through different pathways. This represents a new model of IDR-interactome-dependent transactivation (Fig. 6). Here, we only characterized how particular IDR partners affect gene activation by TFs. It will be of future interest to comprehensively investigate the roles that the newly identified IDR partners play in TF functions besides gene activation, such as chromatin binding location and dynamics, chromatin regulation, and cell phenotype determination. The PhaseID-determined IDR interactomes provide a uniquely valuable resource for understanding the molecular mechanisms underlying transcriptional control through the lens of TF hub formation. Although here we only focus on EWS::FLI1 and Sp1, PhaseID combined with established functional assays will be useful for studying the IDR-mediated functions of numerous other TFs in the future.

**Fig. 6.**
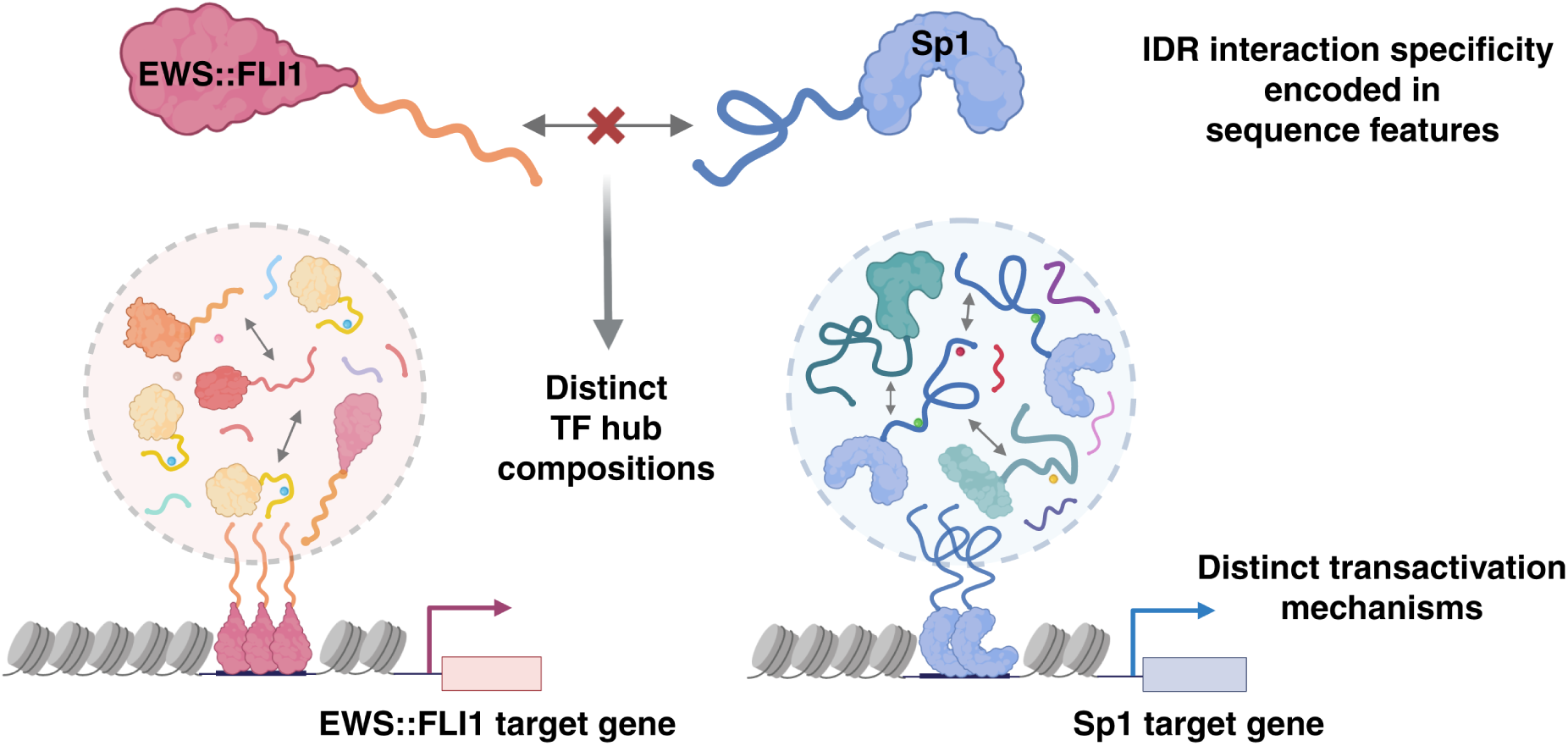
A model for transactivation by different types of transcription factor (TF) hubs. IDR-mediated hubs of EWS::FLI1 and Sp1 feature distinct compositions due to the distinct interactomes of their IDRs, which can lead to different transactivation mechanisms by the two TFs. The selectivity of proteins in binding to specific hubs is encoded in their amino acid sequence features.

An interesting observation we have had from comparing IDR interactomes measured by PhaseID and conventional TurboID-mediated proximity labeling mass spectrometry is that an IDR can alter its interaction selectivity when undergoing LLPS. This is the first time that this property of LLPS has been discovered to our knowledge. The ability of LLPS to tune the interaction selectivity of IDRs may enable a protein to perform new functions by interacting with new partners within its condensates. Consistently, GO enrichment analyses showed that specific biological processes are overrepresented among PhaseID-identified partners of an IDR but not among its TurboID-identified partners, suggesting that proteins containing the IDR may play roles in these processes only upon their LLPS or the formation of related biomolecular assemblies. We speculate that the LLPS dependence of IDR interaction selectivity is due to the IDR’s high local concentration in its condensates, as IDR-mediated protein-protein interactions are often multivalent, i.e., having a variable, concentration-dependent stoichiometry and thus an affinity that increases with protein concentrations (*4*). The concentration dependence of homotypic multivalent interactions has been well characterized (*7*). Here, our findings indicate that the selectivity and affinity of IDR-mediated heterotypic multivalent interactions are likely dependent on protein concentrations as well. Since endogenous proteins often form IDR-mediated biomolecular assemblies with higher local concentrations of the IDR compared to the surrounding cellular environment, even though these concentrations might be lower than those in its LLPS condensates, we speculate that the IDR interactome is at least partially different inside and outside such biomolecular assemblies. It will be of future interest to characterize how an IDR’s interactome may be quantitatively dependent on its concentration and how the concentration dependence may contribute to specific cellular functions. The concentration dependence of IDR interaction selectivity may also explain phenomena such as the requirement for optimal local IDR concentrations for efficient transcriptional activation of endogenous target genes by the IDR-containing TF (*61*, *109*, *114*).

Another finding from analyzing PhaseID-determined IDR interactomes is that the selectivity of proteins in binding to specific IDR condensates is encoded in particular protein sequence features. Intriguingly, PhaseID revealed that proteins in EWS and Sp1 IDR condensates have distinct sequence features, whereas TurboID measurements did not. This suggests that these sequence features play a crucial role in the IDR interaction selectivity of proteins within LLPS condensates, and not in the absence of LLPS. Thus, PhaseID provides a uniquely valuable tool for future studies that decode the sequence grammar underlying the interaction selectivity of numerous IDRs in the context of LLPS or related biomolecular assemblies. Combining PhaseID and the continuously advancing computational tools for protein sequence feature analysis (*93*, *94*, *115*) will pave the way for building a comprehensive sequence-interaction-function paradigm for the dark proteome.

Taken together, PhaseID provides a powerful tool for elucidating both the physicochemical origins and functional impacts of IDR interaction selectivity through the lens of LLPS. This work utilizes PhaseID to uncover the existence of various transactivation pathways. Given the extensive involvement of IDRs and biomolecular condensates in cell biology and pathology, PhaseID is expected to facilitate new discoveries in the study of numerous cellular processes and diseases in the future.

## Supporting information

PhaseID_Supplementary Materials

## Acknowledgments

We thank Jiapei Miao, Michael Xiong, and Jonathan Banh for their participation in the early investigation, Max Staller, Zikun Zhu and William Rosencrans for their helpful discussions, Diana Huynh and Lai Chai Foong for instrument training, Luke Lavis for providing the fluorescent HaloTag ligand, and Dianne Newman and Mitchell Guttman for their critical reading of the manuscript. We also thank the Proteome Exploration Laboratory and Biological Imaging Facility at California Institute of Technology for providing helpful equipment and expertise. Schematics of the model illustrated in Figure 6 were created using BioRender (http://biorender.com).

## Funding

This work is supported by

Shurl and Kay Curci Foundation Research Grant (S.C.)

Pew-Stewart Scholar for Cancer Research Award (S.C.)

Searle Scholar Award (S.C.)

Merkin Innovation Seed Grant (S.C.)

Mallinckrodt Research Grant (S.C.)

Margaret E. Early Medical Research Trust Grants (S.C.)

Alex’s Lemonade Stand Foundation Innovation Grant under award number 1260879 (S.C.)

A grant from the Jonsson Comprehensive Cancer Center (S.C.)

Caltech Beckman Institute Endowment Funds that partially supports the Proteome Exploration Laboratory

## Author contributions

Conceptualization: SC, QH

Methodology: QH, SC, TFC, BQ, TYW

Investigation: QH, YZ, BQ, TYW, TFC, SC

Visualization: QH

Funding acquisition: SC, TFC

Supervision: SC, TFC

Writing – original draft: SC, QH, YZ, BQ, TYW

Writing – review & editing: SC, QH, YZ, TFC, BQ, TYW

## Competing interests

Authors declare that they have no competing interests.

## Data and materials availability

Mass spectrometry data have been deposited to the ProteomeXchange Consortium via the PRIDE partner repository (https://www.ebi.ac.uk/pride/) (*116*) with the dataset identifier PXD070336. All other data are available in the main text or supplementary materials.

## Supplementary Materials

Materials and Methods

Figs. S1 to S12

Tables S1 to S3

References (*120*–*121*)

## Notes

### Competing Interest Statement

The authors have declared no competing interest.

### Summary of Updates

Main text updated to include additional content, such as new discussion on the role of phase separation in tuning disordered protein interactomes and functions; Fig. 1D, 1E, 2D and Table S1 updated for clarity; Fig. 4B, supplementary Fig. S7, S8 and Table S2, S3 updated to include new data; Methods and supplementary figure captions updated for clarity; Supplemental References updated for completeness; Acknowledgments updated.

