## Supplementary material for "Decoding Condensate Composition Reveals Tunable Interactomes and Functions of Disordered Proteins": PhaseID_Supplementary Materials

**This PDF file includes:**

Materials and Methods
Supplemental Figures: Figs. S1 to S12
Tables S1-S3
Supplemental References (*120-121*)

### Materials and Methods

#### Cell culture, stable cell line construction and imaging sample preparation

Human embryonic kidney 293 (HEK293) cells were grown in MEM (Corning, 10-010-CV) supplemented with 10% fetal bovine serum (Cytiva, SH30396.03). Human U2OS osteosarcoma cells containing a LacO array (~50,000 LacO repeats) in the genome, as described in Janicki et al. (*35*) were grown in low glucose DMEM (Thermo Fisher, 10567014) supplemented with 10% FBS and 1% penicillin-streptomycin (Thermo Fisher, 15140122). Human A673 Ewing sarcoma cells (ATCC, CRL-1598), and EWS::FLI1-Halo knock-in A673 cells, as described in Chong et al. (*5*), were grown in high-glucose DMEM (Thermo Fisher, 10566016) supplemented with 10% FBS and 1% penicillin-streptomycin. HEK293, U2OS, and A673 cells were cultured at 37°C with 5% $CO_2$.

Stable HEK293 cell lines expressing PhaseID or TurboID constructs were established using PiggyBac-mediated transposition followed by G418 selection. The cDNAs were cloned into a PiggyBac vector carrying a G418 resistance gene and co-transfected with a SuperPiggyBac transposase plasmid using Lipofectamine 3000 (Thermo Fisher, L3000015) following the manufacturer's instructions. 48 hrs after transfection, cells were selected with 1 mg/mL Geneticin™ G418 (Thermo Fisher, 10131035) until all untransfected control cells died (~1 week). Established lines were maintained in 0.5 mg/mL G418 for an additional week before cryopreservation.

To image live HEK293 cells stably expressing PhaseID constructs, cells were seeded in 35-mm glass-bottom dishes (No. 1.5, 14-mm glass diameter; MatTek, P35G-1.5-14-C) 24 hrs prior to imaging. 500 nM rapalog A/C heterodimerizer (Takara Bio, 635056) was added for the Rapalog (+) condition.

To image fixed cells for the LacO array assay, U2OS cells containing the LacO array were plated onto 18-mm No. 1 circular coverslips (VWR, 48380-046) 48 hrs prior to imaging. Cells were co-transfected 24 hrs before imaging with a plasmid encoding mCherry-labeled protein of interest and a plasmid encoding either EYFP-EWS(2–264)-LacI-NLS or EYFP-LacI-NLS using Lipofectamine 3000. Following transfection, cells were rinsed with PBS (Thermo Fisher, 18912014), fixed in 4% paraformaldehyde (Electron Microscopy Sciences, 15710) in PBS for 15 min, and rinsed three times with PBS for 5 min each. Then the coverslips were mounted upside down in VECTASHIELD antifade mounting medium (Vector Laboratories, H-1000) on Superfrost Plus microscope slides (Fisherbrand, 12-550-15).

To measure the subcellular distribution of biotinylated proteins in HEK293 cells stably expressing PhaseID or TurboID constructs, the cells were seeded onto 18-mm No. 1 circular coverslips 24 hrs prior to treatment. For Rapalog (+) conditions, 500 nM rapalog was added for the indicated incubation times, followed by 50 μM biotin treatment (Sigma-Aldrich, B4501) for 1 hr (PhaseID) or 10 min (TurboID). Cells were then fixed in 4% paraformaldehyde in PBS for 15 min, rinsed three times with PBS for 5 min each, permeabilized with 2.5% Triton X-100 (VWR, M143-1L) for 3 min, washed twice with PBS for 10 min each, and blocked with 5% BSA (Goldbio, A-420-500) for 20 min. Cells were then labeled with AF750-conjugated streptavidin (see Table S1 for details) in PBS containing 0.1% Tween-20 (VWR, M147-1L) and 0.3% BSA for 1 hr at room temperature in the dark, followed by two washes with PBS for 5 min each. Then the coverslips were mounted upside down in VECTASHIELD antifade mounting medium on Superfrost Plus microscope slides.

To evaluate colocalization between endogenous EWS:::FLI1-Halo and PhaseID-identified EWS IDR partners in the knock-in A673 cells, HaloTag labeling and immunostaining were performed prior to imaging. Cells were plated onto 18-mm No. 1 circular coverslips 24 hrs before staining and incubated with 200 nM JFX646 dye (*119*) in the full growth media for A673 cells described above for 1 hr, followed by two rounds of rinsing, each consisting of four washes with PBS and a subsequent 15-min incubation in fresh medium. Fixation and permeabilization were performed as described above. Cells were then incubated with primary antibodies (see Table S1 for details) in PBS containing 0.1% Tween-20 and 0.3% BSA overnight at 4 °C. The cells were washed three times with PBS for 5 min each and incubated with secondary antibody (see Table S1 for details) for 1 hr at room temperature in the dark. The cells were washed three times with PBS for 5 min each, and nuclei were then counterstained with 10 µM Hoechst 33342 (Thermo Fisher, 62249) at room temperature for 10 min in the dark. Finally, the coverslips were mounted upside down in VECTASHIELD antifade mounting medium on Superfrost Plus microscope slides.

### Confocal fluorescence imaging and image analysis

#### Live-cell imaging of HEK293 cells that stably express PhaseID or TurboID constructs

Live-cell confocal fluorescence imaging was performed on a Nikon AX R microscope equipped with a Ti2 inverted base, a Plan Apo λD 60×/1.42 oil-immersion objective, a heated stage, and a full incubation chamber maintaining 37 °C and 5% $CO_2$. To visualize stable expression of PhaseID or TurboID constructs in HEK293 cells and to confirm puncta formation in PhaseID–IDR expressing HEK293 cells upon rapalog induction, images were acquired at 1024 × 1024 pixels (0.2877 µm/pixel) with NIS-Elements AR software. Z-stacks were collected at 1.5-µm intervals to a depth of 46.5 µm (31 sections). EGFP and mCherry were excited with 488-nm and 561-nm lasers, respectively, using identical imaging settings across samples. Time-lapse z-stacks were acquired every hour.

#### LacO array assay imaging and analysis

Confocal fluorescence images were acquired on the Nikon AX R microscope. For a LacO array assay, images were collected at 512 × 512 pixels (0.0575 µm/pixel). Z-stacks were acquired at 0.2-µm intervals to cover the LacO array signal, typically to a depth of ~5 µm (25 sections). EYFP and mCherry were excited using 488-nm and 561-nm lasers, respectively, using identical imaging settings across samples. For visualization, the mCherry channel for the construct mCherry-LIMD1-NLS was processed with Enhance Local Contrast (CLAHE) in Fiji (ImageJ, NIH) (Fig. 2D, Fig. S3C); all quantification was performed on raw, unprocessed images.

For each U2OS cell, the LacO array locus was identified in the EYFP channel by locating a 10 × 10 pixel region with the highest mean EYFP intensity in the maximum-intensity projection image of the z-stack. The center pixel of the identified region represents the center of the LacO array, and the z-slice with the highest mean EYFP intensity within the same region across the z-stack was identified as the focal plane of the LacO array. Radial intensity profiles were extracted from the center of the LacO array in the selected z-slice, and its periphery was defined as the elbow point of the intensity decay curve. The same coordinates were applied to the mCherry channel, and the mCherry enrichment was calculated as the ratio of mean intensity at the LacO center relative to the mean intensity at the periphery. Image preprocessing, periphery detection, and quantification were automated in Python to ensure reproducibility across datasets.

Statistical analyses were performed in Python with results summarized across cells ($n \geqslant 20$). For each POI (protein of interest), we measured the enrichment of mCherry-POI-NLS at the LacO array bound by either EYFP-EWS IDR-LacI-NLS or EYFP-LacI-NLS (control), and compared the mCherry enrichment values using Welch's two-sample *t*-test, with significance thresholds of $p < 0.05$ (*), $p < 0.01$ (**), $p < 0.001$ (***), and $p < 0.0001$ (****), not significant (ns). A significant higher enrichment for the EYFP-EWS IDR-LacI-NLS condition than the control suggests EWS IDR-POI interactions at the LacO array.

<u>Immunofluorescence imaging of A673 cells and analysis</u>

Confocal fluorescence images were acquired on the Nikon AX R microscope. For immunofluorescence in the knock-in A673 cells, images were collected at 512 × 512 pixels (0.0575 μm/pixel). Z-stacks were acquired at 0.3-μm intervals to cover the EWS::FLI1 signal, typically to a depth of ~6 μm (20 sections). Hoechst 33342, AF488, and JFX646 were excited with 405-nm, 488-nm, and 640-nm lasers, respectively, using identical imaging settings across samples. To reduce detector noise, images in Fig. S1B were processed using the "Despeckle" function in Fiji. "Despeckle" applies a 3 × 3 median filter that replaces each pixel with the median value of its local neighborhood.

To evaluate the enrichment of POI immunostained by AF488-labeled antibody at JFX646-labeled endogenous EWS::FLI1-Halo hubs, AF488 and JFX646 channels of confocal stacks were analyzed with a custom Python workflow. For each cell, a JFX646 maximum-intensity projection was generated, and nuclear masks were derived from the two-channel projection by Gaussian filtering, Otsu thresholding, and binary erosion to restrict analysis to nuclei. EWS::FLI1-Halo hub regions were segmented in the projection image of the JFX646 channel by applying a Difference-of-Gaussians (DoG) filter, followed by intensity thresholding and connected-component analysis to identify contiguous high-intensity blobs. To obtain precise coordinates of individual hubs, non-maximum suppression within each blob was applied to identify the pixel with the maximal DoG-filtered intensity, which was then defined as the x–y coordinate of the hub locus. Background, non-hub loci in the nucleus were identified by analyzing a dark DoG map outside of hub regions, where the inverted Gaussian scales highlighted local intensity minima. Using intensity thresholding and connected-component analysis, contiguous dark blobs were segmented from the dark DoG map. For each blob, the pixel with the maximal dark DoG-filtered intensity was selected as the x–y coordinate of the background locus, with non-maximum suppression ensuring that only one locus was retained per dark region. To determine the focal plane for each hub or background locus, the z-slice with the maximum mean JFX646 intensity across a 5 × 5 pixel window centering the previously defined x-y coordinate was selected. The same focal plane and x–y coordinates identified from the JFX646 channel were then applied in the 488 channel to ensure matched sampling. Annotated overlays of hubs (red) and backgrounds (cyan) were generated on EWS::FLI1-Halo channel maximum projections (Fig. S4B), and corresponding statistics were recorded for manual review.

Centering each hub or background location, a 61 × 61 pixel square ROI was extracted from both channels at the selected z-slice. Hub ROIs and background ROIs were then averaged separately to generate mean hub and mean background images. For each immunofluorescence sample, 10–20 cells were analyzed, resulting in a total of 1,400–2,500 EWS::FLI1 hubs, along with an equal number of background regions processed.

For each sample, background-free two-color images (Fig. 3B) were obtained by subtracting the mean background images of each channel from its corresponding mean hub images. Horizontal intensity profiles were extracted from a centered 18 × 18 pixel (≈approximately 1 μm × 1 μm) square region of each channel by averaging the pixel intensities along the columns. For each channel, profiles were background-corrected by subtracting a baseline estimated from the low-intensity flanking areas, and then normalized to the peak intensity.

Imaging AF750-streptavidin-stained HEK293 cells and analysis

Confocal fluorescence imaging was performed on a Leica Stellaris 8 microscope (DMI8-CS inverted base) equipped with a HC PL APO CS2 63×/1.40 oil-immersion objective. For the modified HEK293 cells stained with AF750-streptavidin, images were acquired at 512 × 512 pixels (0.0722 µm/pixel) using LAS X software. Z-stacks were collected at ~0.3-µm intervals to cover the nucleus signal, typically to a depth of ~8.2 µm (25 sections). Excitation was provided by a 405-nm diode laser and a white-light laser (WLL). Images were acquired using a 405-nm laser for Hoechst (not shown), 473-nm excitation for EGFP, 587-nm excitation for mCherry, and 688-nm excitation for AF750-streptavidin. All channels were acquired sequentially to avoid spectral crosstalk. The AF750-streptavidin channel was averaged to improve the signal-to-noise ratio.

Prior to analysis, channels were spatially aligned using a translation-only alignment in Fiji to match the nuclear boundary across channels. Line profiles were generated in Fiji to assess the co-distribution of EGFP-core, mCherry-bait, and AF750-streptavidin signals. Straight ROIs (~4.48 µm) were drawn at identical spatial locations across three channels, and pixel intensities were sampled at native resolution. Profiles were background-corrected and then normalized to the global maximum across all channels and conditions to enable direct comparison of signal overlap.

Confocal images were further analyzed using a custom Python pipeline for automated nucleus segmentation, puncta detection, and size quantification. All analyses were performed on maximum-intensity projections of z-stacks. Nuclei were segmented from the AF750-streptavidin channel using Gaussian smoothing, Otsu thresholding, and a marker-controlled watershed algorithm. Biotinylated puncta were identified within segmented nuclei in the AF750-streptavidin channel by bandpass filtering and local maxima detection, followed by signal-to-noise and z-score-based filtering to avoid background mis-selection. Puncta diameter was computed from radial profiles using two complementary criteria: a half-maximum threshold and an energy-based radius enclosing 70% of the total intensity, and the smaller value was reported. Per-nucleus puncta numbers, and diameter distributions were computed after excluding sub-threshold objects (< 3 px). Segmentation and puncta overlays were manually reviewed for quality control (Fig. S1C).

General acquisition settings

Before image acquisition, laser power and detector gain were adjusted to avoid pixel saturation. For multi-color imaging, appropriate emission detection windows were applied, and the absence of spectral bleed-through was validated using single-labeled control samples imaged under identical acquisition settings. The pinhole was set to 1 Airy unit at the chosen objective settings. Unless otherwise specified, image display adjustments (brightness/contrast and color assignment) and image export were performed in Fiji.

**siRNA knockdown, RT-qPCR and analysis**

A673 cells were seeded onto 18-mm No. 1 circular coverslips 24 hrs before transfection. Cells were transfected with 10 nM Silencer Select siRNA (Thermo Fisher, Table S2) using Lipofectamine RNAiMAX (Thermo Fisher, 13778150) following the manufacturer’s instructions and incubated for 72 hrs before downstream processing. Total RNA was extracted using TRIzol reagent (Thermo Fisher, 15596026) following the manufacturer's instructions. Equal amounts of total RNA from each sample were reverse transcribed into cDNA using the PrimeScript RT Master Mix (Takara Bio, RR036A). The resulting cDNA was diluted and subjected to real-time quantitative PCR using PowerUp SYBR Green Master Mix (Applied Biosystems, A25776) on a QuantStudio 5 Real-Time PCR System (Applied Biosystems). Ct values from each reaction were collected, and each sample was measured in four technical replicates to ensure measurement reproducibility. Relative mRNA expression levels were calculated using the ΔΔCt method. Data are presented as $\log_2$(fold change) relative to control samples (Fig. 4B), and statistical significance was determined using Welch's two-sample *t*-test, with significance thresholds of $p < 0.05$ (*), $p < 0.01$ (**). Primer sequences for each target gene are listed in Table S3.

Knockdown efficiency was verified by either Western blot or quantitative PCR, as detailed in Supplementary Figs. S7 and S8. Only siRNAs that achieved robust knockdown and showed no technical issues in the RT–qPCR assay, as listed in Table S2, were used for quantifying subsequent EWS::FLI1 and Sp1 target gene responses (Fig. 4B).

For quantification of target gene responses upon siRNA treatment, each data point represents the averaged effect of one or multiple siRNAs targeting a given interactor on the transcription level of a specific target gene. For Sp1 and EWS knockdown experiments, transcription levels were assessed across four validated target genes, respectively (EWS::FLI1*: CYP4F22*, *CCK*, *PPP1R1A*, and *NR0B1*; Sp1: *BIRC5*, *B4GALT5*, *MMP14*, and *IGF1R*).

### Western blot

For PhaseID cell lines, lysates were prepared to verify biotinylation of the EGFP-core and mCherry-IDR bait upon rapalog induction, and to quantify rapalog-dependent changes in biotinylation. Cells were lysed in EasyPep lysis buffer (Thermo Fisher, A45735) supplemented with 25 U/mL Turbo DNase (Thermo Fisher, AM2238) and 1× protease inhibitor cocktail (Thermo Fisher, A32963). Lysates were clarified by centrifugation at 15,000 × g for 15 min at 4 °C. Protein concentrations were determined using Pierce BCA Protein Assay Kit (Thermo Fisher, 23227). Proteins were mixed with 4× Laemmli sample buffer (Bio-Rad, 1610747), boiled at 95 °C for 10 min, and briefly centrifuged prior to SDS–PAGE. A total of 10 µg of protein per lane was loaded on 4–20% Mini-PROTEAN TGX Precast Protein Gels (Bio-Rad, 4561096) alongside 3 µL Precision Plus Protein Kaleidoscope Prestained Protein Standards (Bio-Rad, 1610375). Proteins were transferred to nitrocellulose membranes (Cytiva, 10600041) using the Trans-Blot Turbo Transfer System (Bio-Rad) with the mini gel cycle for 7 min. Membranes were stained with 1× Ponceau S solution (Thermo Fisher, A40000279) for 10 min to confirm equal loading, imaged on a ChemiDoc imaging system (Bio-Rad), and then destained with Milli-Q water for 15 min. Membranes were blocked in 3% BSA in TBST for 60 min at room temperature, and then incubated overnight at 4°C with primary antibodies (see Table S1 for details) diluted in 3% BSA in TBST. After three washes in TBST for 5 min each, membranes were sequentially incubated with secondary antibodies (see Table S1 for details) and IRDye 800CW Streptavidin, each for 60 min at room temperature in 3% BSA in TBST. Membranes were washed three times in TBST for 5 min each and rinsed once in Mili-Q water before imaging. Blots were then imaged using the Odyssey imaging system (Licor).

For siRNA knockdown verification, cells were lysed in RIPA lysis buffer (Thermo Fisher, 89900) supplemented with 25 U/mL Turbo DNase and 1× protease inhibitor cocktail. Lysates were clarified by centrifugation at 15,000 × g for 5 min at 4 °C, and protein concentrations were determined using the Bradford assay (Bio-Rad, 5000006). The resulting supernatants were subjected to SDS-PAGE followed by immunoblotting as described previously.

### Proximity labeling and mass spectrometry

Cells cultured in 15-cm dishes were treated with 50 µM biotin in the culture medium for 1 hr (PhaseID cell lines) or 10 min (TurboID cell lines). Cells were then rinsed with cold PBS, detached using TrypLE Select (Thermo Fisher, 12563011), and pelleted by centrifugation at 3,000 × g for 4 min at 4 °C in low protein-binding tubes (Thermo Fisher, 90410). Cell pellets were lysed in EasyPep lysis buffer supplemented with Pierce Universal Nuclease (Thermo Fisher, 88700). Lysates were sonicated at 15 amps for 5 s × 3 cycles with 5 min pause on ice between cycles, and clarified by centrifugation at 16,600 × g for 15 min at 4 °C. Protein concentration was determined using the Pierce BCA Protein Assay Kit.

A total of 5 mg of clarified lysate was incubated with 50 µL of washed Pierce streptavidin magnetic beads (Thermo Fisher, 88817) for 1 hr at 4 °C with gentle agitation. After binding, the beads are washed five times with EasyPep lysis buffer and twice with 50 mM HEPES (pH 8.0; Fisher Scientific, BP310-100). On-bead digestion was performed in 8 M urea (Thermo Fisher, 29700) containing 5 mM TCEP-HCl (Thermo Fisher, 20490) for 20 min at 37 °C with shaking at 750 rpm, followed by alkylation with 15 mM 2-chloroacetamide (MP Biomedicals, 15495580) for 15 min at 37 °C with shaking at 750 rpm. Beads were then incubated with 200 ng Lys-C endoproteinase (Thermo Fisher, 90051) in the same buffer for 4 hrs at 37 °C with shaking at 750 rpm. The mixture was subsequently diluted to 2 M urea, supplemented with 1 mM $CaCl_2$, and digested with 300 ng trypsin (Thermo Fisher, 90057) in 50 mM HEPES overnight at 37 °C with shaking at 750 rpm. After digestion, the supernatants were acidified with 0.6% trifluoroacetic acid (TFA; Thermo Fisher, 85183). C18 spin columns (Thermo Fisher, 89870) were pre-equilibrated by washing twice with 50% acetonitrile (ACN; Fisher Scientific, A955-1), followed by two washes with 0.5% TFA in 5% ACN.

Acidified peptide mixtures were loaded onto the columns and washed with 0.5% TFA in 5% ACN. Bound peptides were eluted twice with elution buffer containing 70% ACN and 0.2% formic acid (FA; Thermo Fisher, 85178), and the eluates were vacuum-dried (Eppendorf, Vacufuge Plus) for 4 h and stored at –80 °C prior to LC-MS/MS analysis on an Orbitrap Eclipse Tribrid mass spectrometer (Thermo Fisher).

Dried peptides were reconstituted in 20 µL of 0.1% FA (Thermo Fisher, A117-50), and 4–8 µL were injected for LC-MS/MS analysis on an Orbitrap Eclipse Tribrid mass spectrometer equipped with a Vanquish Neo UHPLC system. Peptides were separated on an Aurora C18 column (25 cm × 75 µm, 1.7 µm C18; Ion Opticks, AUR3-25075C18-TS) at a flow rate of 0.35 µL/min and ionized at 1.8 kV in the positive ion mode. The active gradient was composed of 6% B (3.5 min), 6–25% B (41.5 min), 25–40% B (15 min), 40–98% B (2 min), and 98% B (5 min). Mobile phase A consisted of 2% ACN and 0.2% FA in water (Fisher Scientific, W6212); mobile phase B consisted of 80% ACN and 0.2% FA in water. MS1 spectra were acquired in the Orbitrap at the resolution of 120,000 from 375 to 1,600 m/z with automatic gain control (AGC) target of 250% and a maximum injection time of 50 ms. MS2 spectra were acquired in the ion trap using fast scan mode with quadrupole isolation (1.2 m/z window), higher-energy collisional dissociation (HCD, 30%). Only precursors with charge states of +2 to +7 were selected, with dynamic exclusion of 15 s and precursor mass tolerance of 5 ppm. The ion transfer tube was set to 300 °C and the S-lens RF level to 30.

Raw data were processed in Proteome Discoverer (v2.5, Thermo Scientific) using the SEQUEST HT search engine against the Swiss-Prot human database (UniProt 2022, 20,309 entries). Trypsin/P was specified as the digestion enzyme, allowing a maximum of two missed cleavages. Dynamic modifications were set to oxidation on methionine (M, +15.995 Da), deamidation on asparagine (N,Q +0.984 Da) and protein N-terminal acetylation (+42.011 Da). Carbamidomethylation on cysteine (C, +57.021 Da) was set as a fixed modification. The maximum parental mass error was set to 10 ppm, and the MS2 mass tolerance was set to 0.6 Da. The false discovery rate (FDR) was set strictly to 0.01 using the Percolator Node validated by q-value. The relative abundance of parental peptides was calculated by integration of the area under the curve of the MS1 peaks using the Minora LFQ node. The mass tolerance used to align features across runs was set to 5 ppm in the Feature Mapper node.

Mass spectrometry data have been deposited to the ProteomeXchange Consortium via the PRIDE (*116*) partner repository with the dataset identifier PXD070336.

Protein abundances were extracted based on LFQ intensity values and filtered to retain only proteins with ≥2 unique peptides and high-confidence identifications. For downstream analysis, proteins with missing values were imputed using KNN (k=5), followed by median normalization across all replicates.

Differential enrichment analysis was performed by comparing Rapalog (+) and Rapalog (-) conditions across biological replicates using an unpaired Student's *t*-test. For each PhaseID construct, protein abundance was averaged across biological replicates within each condition to obtain a mean abundance for each protein in that condition. $Log_2$(fold change) of Rapalog (+) condition relative to Rapalog (-) condition was then calculated for each protein using the mean abundance values. The $p$ values were converted to $-log_{10}$ scale for visualization. Proteins with $log_2$(fold change) > 0.585 or < –0.585 and $p < 0.05$ were considered significantly enriched in or depleted from the induced condensates, respectively. To identify IDR-dependent interactions, proteins that also showed similar enrichment or depletion behavior in the corresponding no IDR control PhaseID dataset ($p < 0.05$ and $log_2$(fold change) > 0.585 or < –0.585) were excluded from the IDR PhaseID dataset to generate the IDR interactome. The volcano plots were generated using VolcaNoseR (*120*).

For each TurboID construct, differential enrichment analysis was performed by comparing the IDR and no IDR conditions across biological replicates using the same unpaired Student's *t*-test described above. Protein abundance was first averaged across replicates within each condition to obtain a mean abundance for each protein. Subsequently, $log_2$(fold change) of the IDR condition relative to the no IDR control condition was calculated for each protein using the mean abundances. The $p$ values were transformed to $-log_{10}$ for visualization. Proteins with $log_2$(fold change) > 0.585 and $p < 0.05$ were considered as IDR-dependent interaction partners identified by TurboID.

### GO analysis

Gene Ontology (GO) enrichment was performed using the Functional Annotation Clustering tool from DAVID Bioinformatics Resources (v6.8) (*63*). Proteins identified in each IDR's PhaseID- or TurboID-identified interactome with $p < 0.05$ and $log_2$(fold change) > 0.585 were used as input, with the full Homo sapiens proteome as background. Analyses were conducted under default parameters, using the annotation categories GOTERM_BP_ALL and GOTERM_CC_ALL. Redundant or conceptually overlapping GO terms were manually grouped into broader biological categories. Representative terms were selected for display based on cluster significance and relevance (Fig. 4D). GO terms shown in Fig. S11 were selected based on FDR values (FDR < 0.05) as calculated by the DAVID Bioinformatics Resources.

### Protein–protein interaction network visualization

Protein interaction networks were generated for the subset of PhaseID-identified proteins annotated with transcription-related GO terms, using the STRING database (v12.0) restricted to experimentally supported interactions with a high-confidence score (≥ 0.7). Network clustering was performed using the Markov Cluster Algorithm (MCL) with an inflation parameter of 3 to identify functional modules. The clustering results were manually reviewed, and nodes were annotated based on gene functions. Networks were visualized in Cytoscape (v3.10.3) (*121*).

**Protein sequence analysis**

A total of 140 sequence parameters were computed, comprising 42 global parameters (e.g., FCR, κ, and δ) and 11 local parameters (e.g., linear FCR and linear NCPR) calculated with a blob length of 5 and step size of 1 using localCIDER v0.1.20 (*94*). The blob length defines the window size used to calculate the local sequence parameters. Each local parameter was then calculated per protein sequence by its maximum, mean, median, and standard deviation.

Patch regions were defined as contiguous runs of three or more identical or chemically similar residues. If two such runs were separated by fewer than two intervening residues, they were considered a single patch. Only patches of length ≥ 5 amino acids were retained for downstream analyses. For each amino acid or residue class of interest (e.g., negatively charged D/E, positively charged K/R, charged D/E/K/R, polar Q/N/S/T/G/H/C, aliphatic A/L/M/I/V, aromatic F/Y/W, and P), maximum patch length and average patch fraction were calculated, yielding 54 patch-based parameters.

Sequence parameters of proteins in the PhaseID- or TurboID-identified interactome of each IDR were compared with those of the background proteome detected in our mass-spectrometry dataset. For each sequence parameter, values of individual proteins in the IDR-enriched or depleted protein group were normalized to z-scores using the mean and standard deviation of the mass-spectrometry-detected proteome. Each z-score represents how far a protein's parameter value deviates from the proteome average. For each sequence parameter, z-scores of each protein group were compared to those of the proteome using Welch's two-sample *t*-test. Significance was visualized as a signed $-\log_{10}(p$ values), in which the sign reflects the direction of the mean z-score difference: positive when the mean z-score in the protein group is higher than that of the proteome, and negative when it is lower. In Figs. 5 and S12, we highlight a selected subset of parameters that show significant differences ($p < 0.05$) between IDR interactors and the mass-spectrometry-detected proteome.

**Quantification and statistical analysis**

All analyses were performed in Python unless otherwise stated. For mass spectrometry data processing, statistical significance was assessed using an unpaired Student's *t*-test, assuming equal variances between groups. For all other quantitative analyses, Welch's two-sample *t*-test was used, which does not assume equal variances. Unless specified, all tests were two-sided, and nominal *p* values were reported. Significance thresholds were indicated by asterisks as $p < 0.05$ (*), $p < 0.01$ (**), $p < 0.001$ (***), and $p < 0.0001$ (****), not significant (ns).

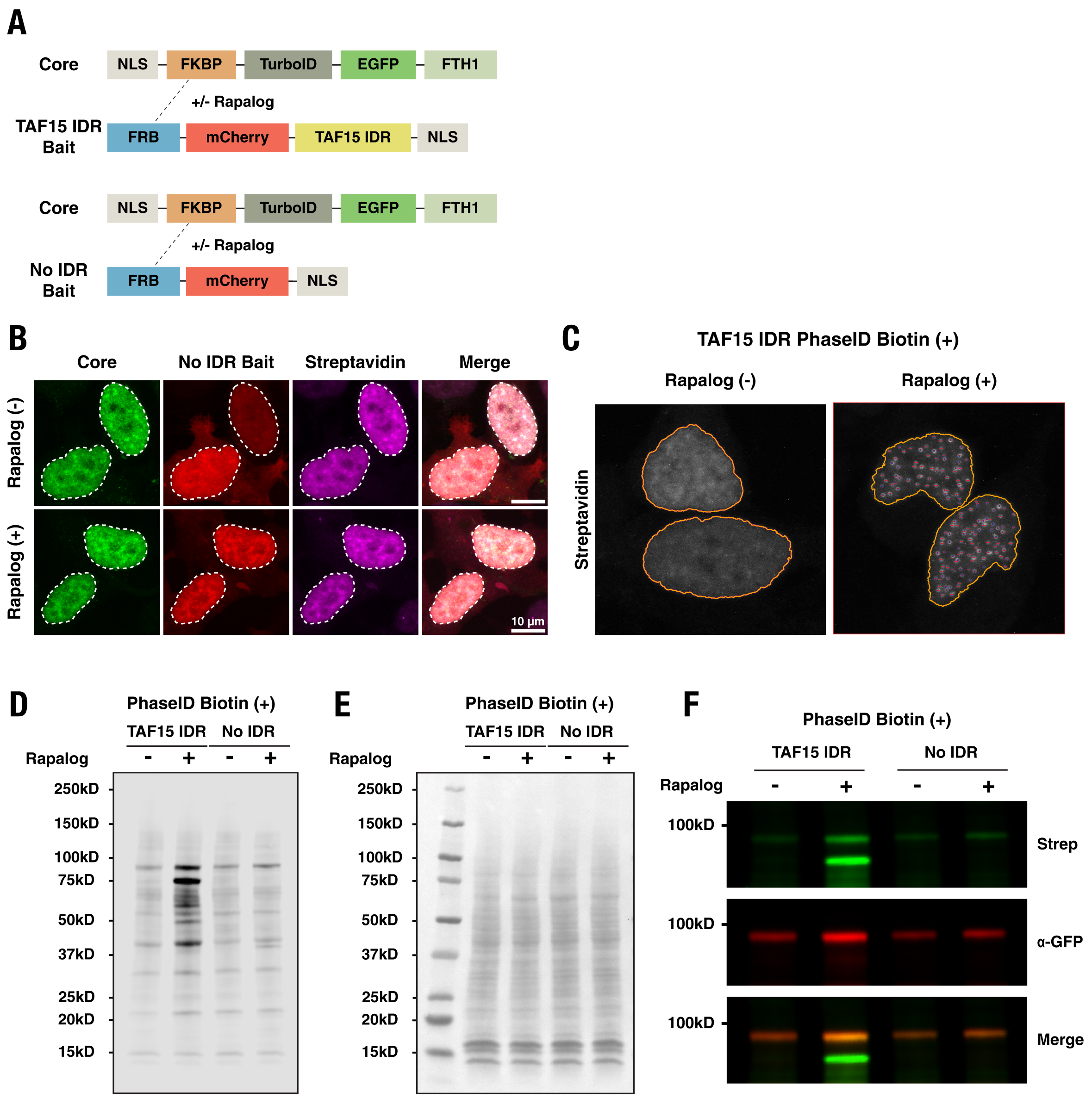


**Fig. S1. Additional information for the design and characterization of the PhaseID system.** (A) Schematics of core and bait constructs for TAF15 IDR PhaseID and no IDR control PhaseID. (B) Confocal fluorescence images of modified HEK293 cells expressing core (green) and no-IDR bait (red) before and after rapalog treatment (500 nM, 10.5 hrs). After the cells were treated with biotin (50 µM, 1 hr), cell staining with AF750-labeled streptavidin (magenta) shows that the biotinylated proteins are distributed nearly homogeneously with no detectable puncta throughout the nuclei. A white dashed contour outlines the cell nucleus. Scale bars, 10 µm.
(C) Representative segmentation results from the automated image-analysis pipeline identifying rapalog-induced TAF15 IDR condensates. Dark blue dots mark the detected puncta, red lines indicate puncta diameters, and yellow outlines define nuclear boundaries.

(D-E) Streptavidin Western blot (D) showing a significant increase in the overall biotinylation levels upon rapalog-induced LLPS of TAF15 IDR in HEK293 cells. In contrast, the modified HEK293 cells for no IDR control PhaseID exhibit minimal change in the overall biotinylation levels after the same rapalog treatment (500 nM, 10.5 hrs). We treated the cells regardless of rapalog treatment with 50 μM biotin for 1 hr. All lanes were loaded with equal amounts of total protein according to Ponceau S staining of the same membrane (E).
(F) Streptavidin (Strep) and anti-GFP (α-GFP) blots demonstrating consistent biotinylation of the EGFP-labeled core (~97.9 kDa) across all conditions. This is expected, as TurboID is part of the core. This result confirms successful TurboID-mediated proximity labeling in the PhaseID system. Equal amounts of total protein were loaded in all lanes, as verified by Ponceau S staining of the same membrane.

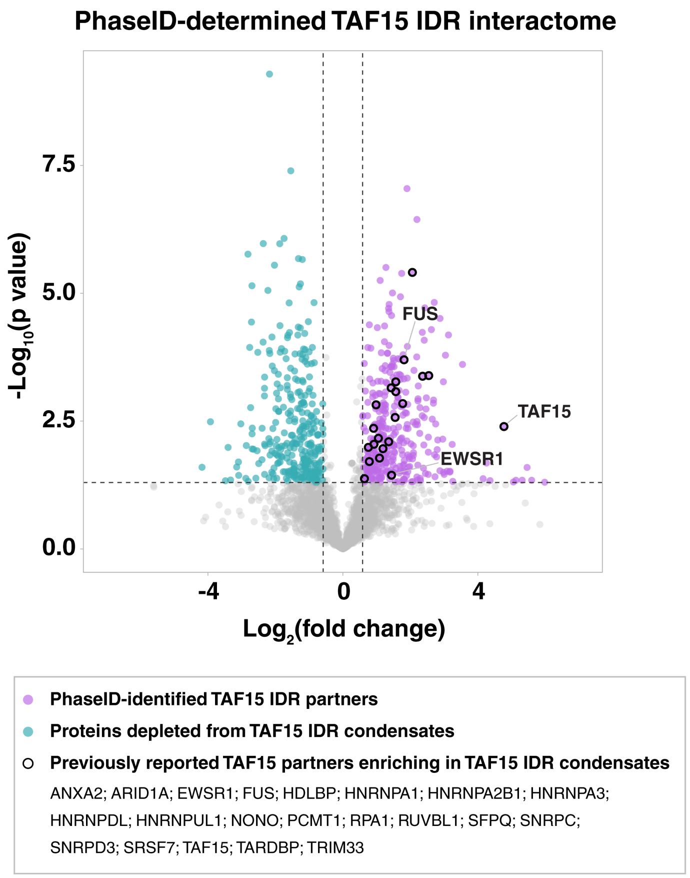


**Fig. S2. Interactome of TAF15 IDR in induced condensates identified by PhaseID.**
Volcano plot showing the PhaseID-determined TAF15 IDR interactome, based on six biological replicates per condition (before and after rapalog treatment). Each protein is represented by a dot in the plot. Proteins significantly enriched in and depleted from the rapalog-induced TAF15 IDR condensates are shown in magenta and blue, respectively. Previously reported TAF15 partners are highlighted with black circles (*117*). The x-axis represents the fold change of the detected protein abundance upon rapalog induction compared with no rapalog condition. The y-axis represents the significance of the fold change. The dashed horizontal line shows the threshold *p* value of statistical significance (0.05). Statistical significance was determined by an unpaired Student's *t*-test. The two dashed vertical lines show the thresholds of abundance fold change (1.5 or 0.67).

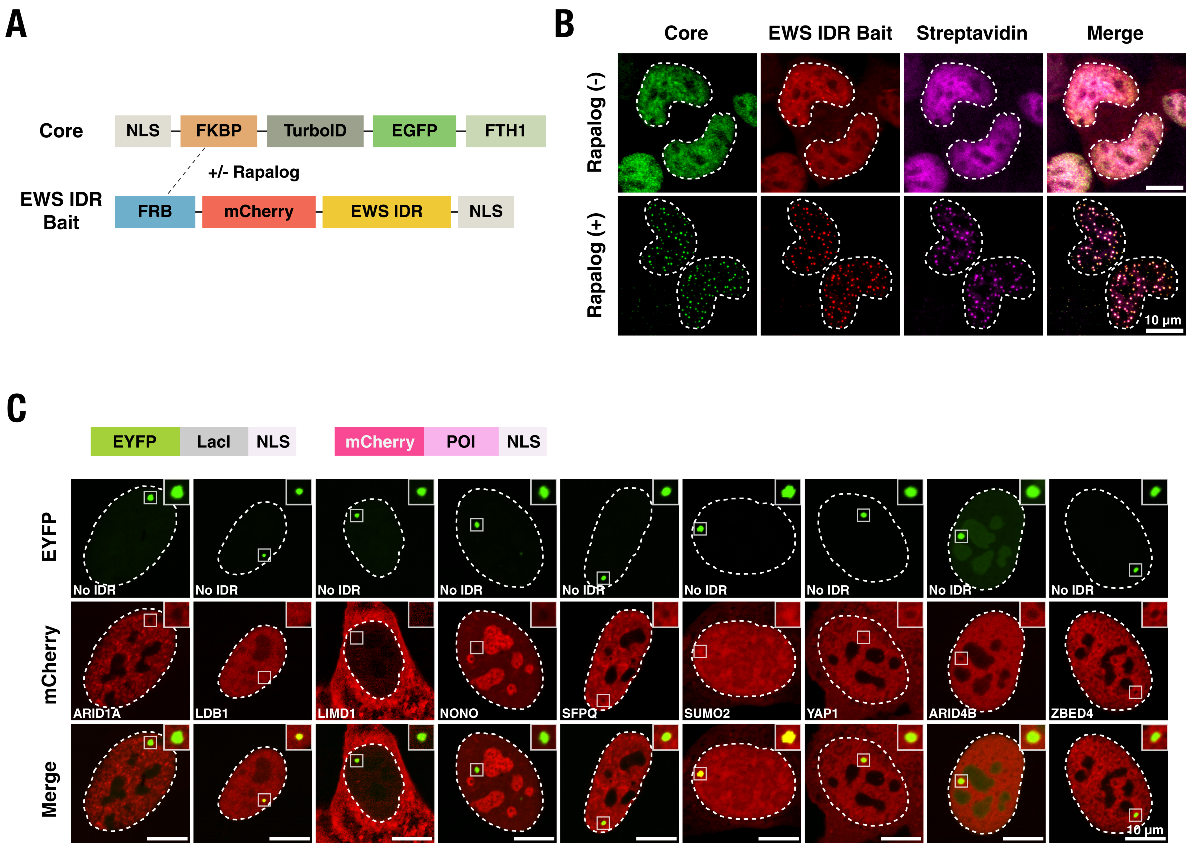


**Fig. S3. Additional information for EWS IDR PhaseID and result validation.**
(A) Schematics of core and bait constructs for EWS IDR PhaseID.
(B) Confocal fluorescence images of modified HEK293 cells expressing core (green) and EWS IDR bait (red) before and after rapalog treatment (500 nM, 2 hrs). After the cells were treated with biotin (50 µM, 1 hr), cell staining with AF750-labeled streptavidin (magenta) shows that protein biotinylation predominantly occurs in the induced EWS IDR condensates. A white dashed contour outlines the cell nucleus. Scale bars, 10 µm.
(C) Confocal fluorescence images of LacO-array-containing U2OS cells co-expressing EYFP-LacI and mCherry-labeled proteins identified by PhaseID. The region surrounding the LacO array is zoomed in. Scale bars, 10 µm. These imaging experiments serve as a control for the LacO array assay (Fig. 2C and D). Quantification of mCherry-POI enrichment at the LacO array bound by EYFP-LacI is provided in Fig. 2E.

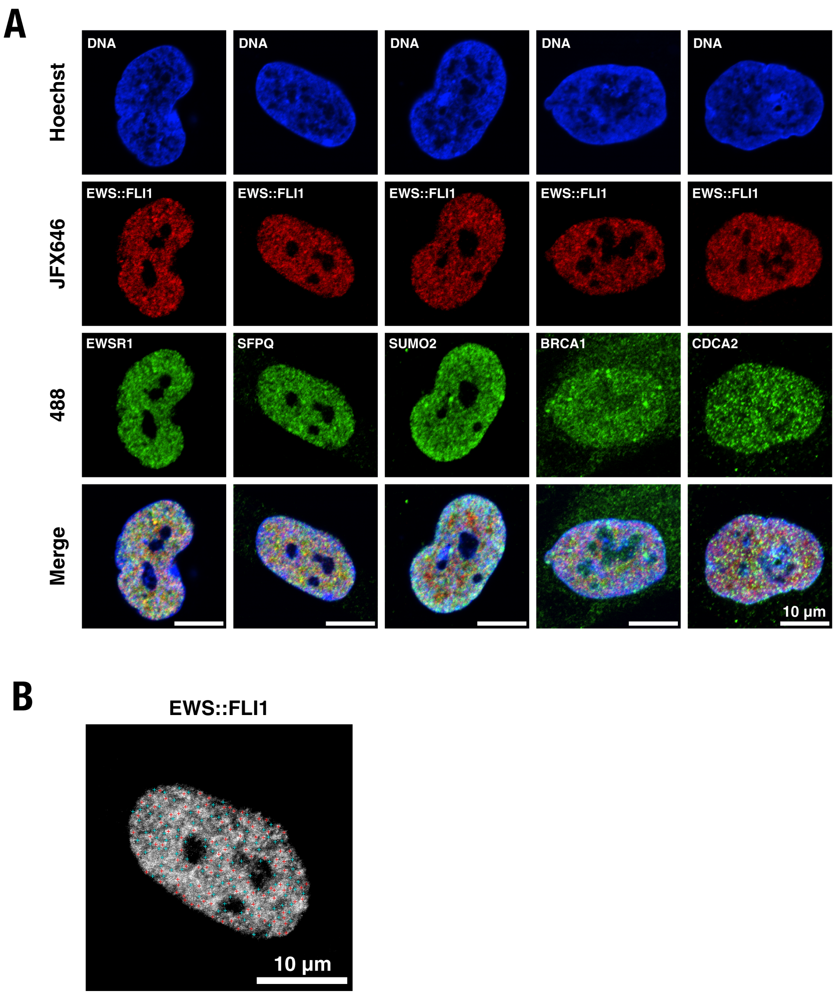


**Fig. S4. Additional information for detecting the PhaseID-identified EWS IDR partners in endogenous EWS::FLI1 hubs.**
(A) Confocal fluorescence images of knock-in A673 cell nuclei showing DNA stained with Hoechst (blue), endogenous EWS::FLI1-Halo labeled with the HaloTag ligand JFX646 (red), and endogenous proteins of interest that are each immunostained with a primary antibody and an AF488-labeled secondary antibody (green). Scale bars, 10 µm. The proteins of interest include three EWS IDR partners (EWSR1, SFPQ, and SUMO2) and two proteins depleted from the induced EWS IDR condensates (BRCA1 and CDCA2). These images were used to generate the averaged two-color fluorescence images shown in Fig. 3B.
(B) Representative image of a knock-in A673 cell nucleus showing the automated selection of fluorescence foci in the EWS::FLI1-Halo channel. Red crosses mark the detected EWS::FLI1 hubs, and blue crosses mark the background sampling points within the nuclear boundary, which were used for subtracting the background intensity to generate Fig. 3B. Scale bar, 10 µm.

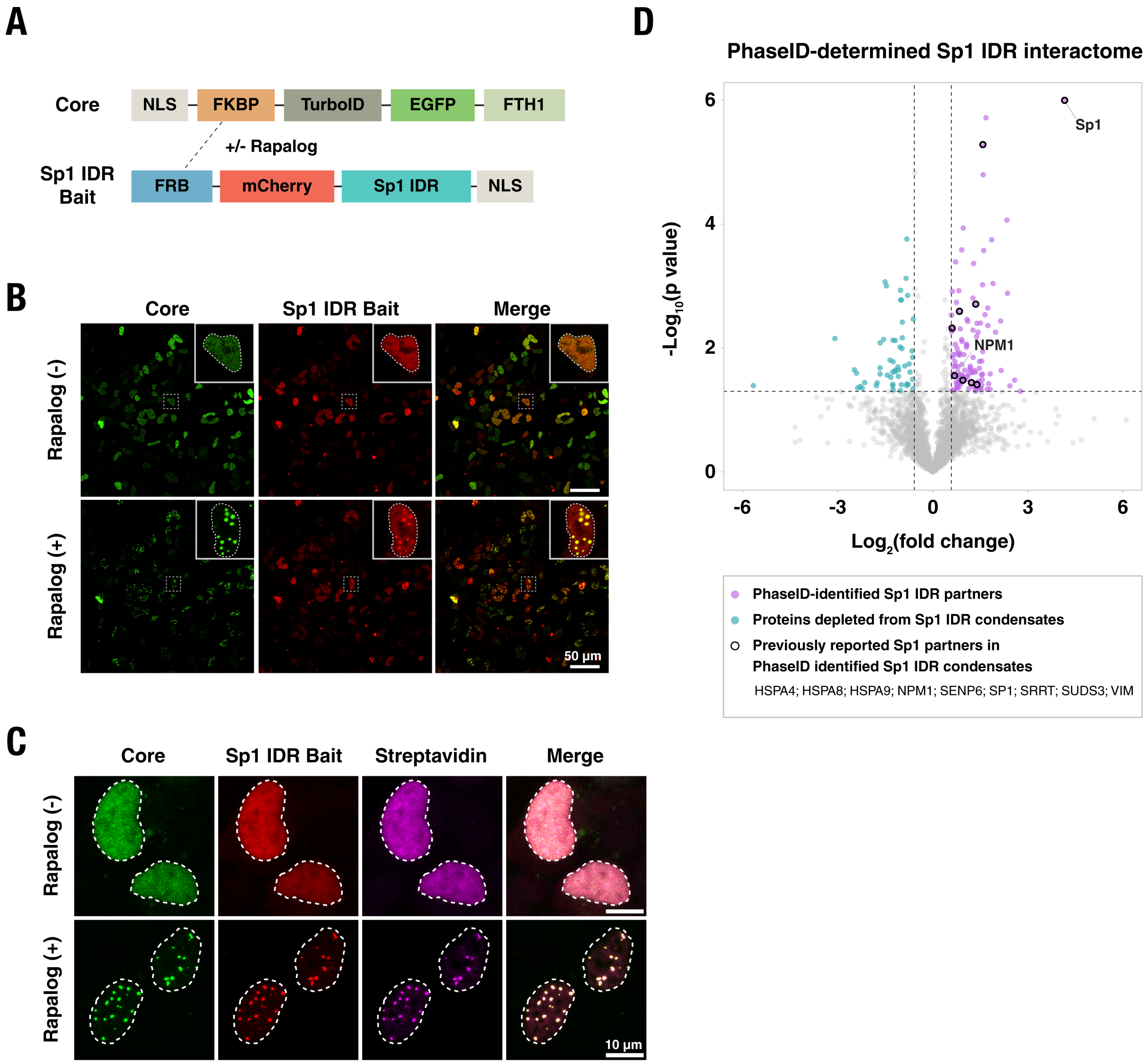


**Fig. S5. PhaseID of Sp1 IDR.**
(A) Schematics of core and bait constructs for Sp1 IDR PhaseID.
(B) Confocal fluorescence images of modified HEK293 cells expressing the Sp1 IDR PhaseID constructs before and after rapalog treatment (500 nM, 2 hrs). Rapalog induces LLPS of Sp1 IDR in live cells. The distributions of core and bait are visualized in EGFP (green) and mCherry (red) channels, respectively. A representative cell under each condition is zoomed in. A white dashed contour outlines the cell nucleus. Scale bar, 50 μm.
(C) Confocal fluorescence images of modified HEK293 cells expressing core (green) and Sp1 IDR bait (red) before and after rapalog treatment (500 nM, 2 hrs). After the cells were treated with biotin (50 μM, 1 hr), cell staining with AF750-labeled streptavidin (magenta) shows that protein biotinylation predominantly occurs in the induced EWS IDR condensates. A white dashed contour outlines the cell nucleus. Scale bars, 10 μm.

(D) Volcano plot showing the PhaseID-determined Sp1 IDR interactome, based on five biological replicates per condition (before and after rapalog treatment). Each protein is represented by a dot in the plot. Proteins significantly enriched in and depleted from the rapalog-induced Sp1 IDR condensates are shown in magenta and blue, respectively. Previously reported Sp1 partners are highlighted with black circles (*117*). The x-axis represents the fold change of the detected protein abundance upon rapalog induction compared with no rapalog condition. The y-axis represents the significance of the fold change. The dashed horizontal line shows the threshold $p$ value of statistical significance (0.05). Statistical significance was determined by an unpaired Student's $t$-test. The two dashed vertical lines show the thresholds of abundance fold change (1.5 or 0.67).

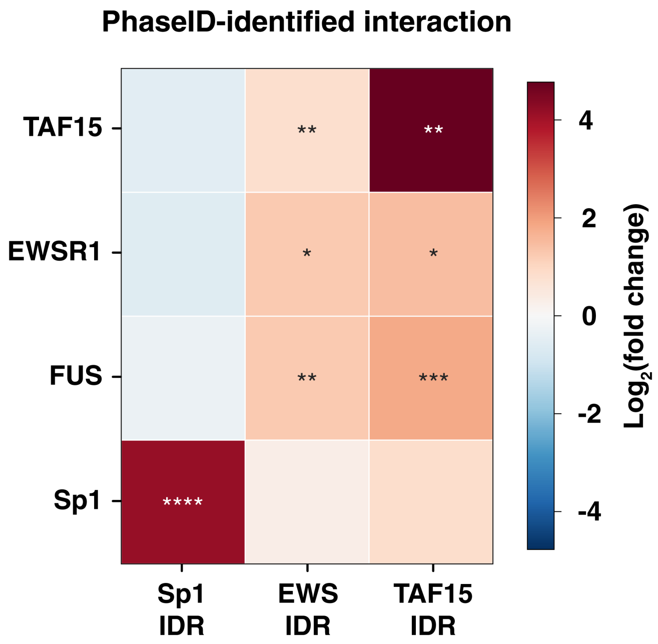


**Fig. S6. PhaseID identifies interactions of FET family proteins and Sp1 in different IDR condensates.**

Heatmap showing the enrichment ($\log_2$(fold change)) of FET family proteins (TAF15, EWSR1, and FUS) and Sp1 in the induced condensates of the indicated IDR (TAF15 IDR, EWS IDR, or Sp1 IDR). Cells are colored based on the $\log_2$(fold change) values. Asterisks denote the statistical significance levels determined by PhaseID: $p < 0.05$ (*), $p < 0.01$ (**), $p < 0.001$ (***), $p < 0.0001$ (****); Unpaired Student's *t*-test.

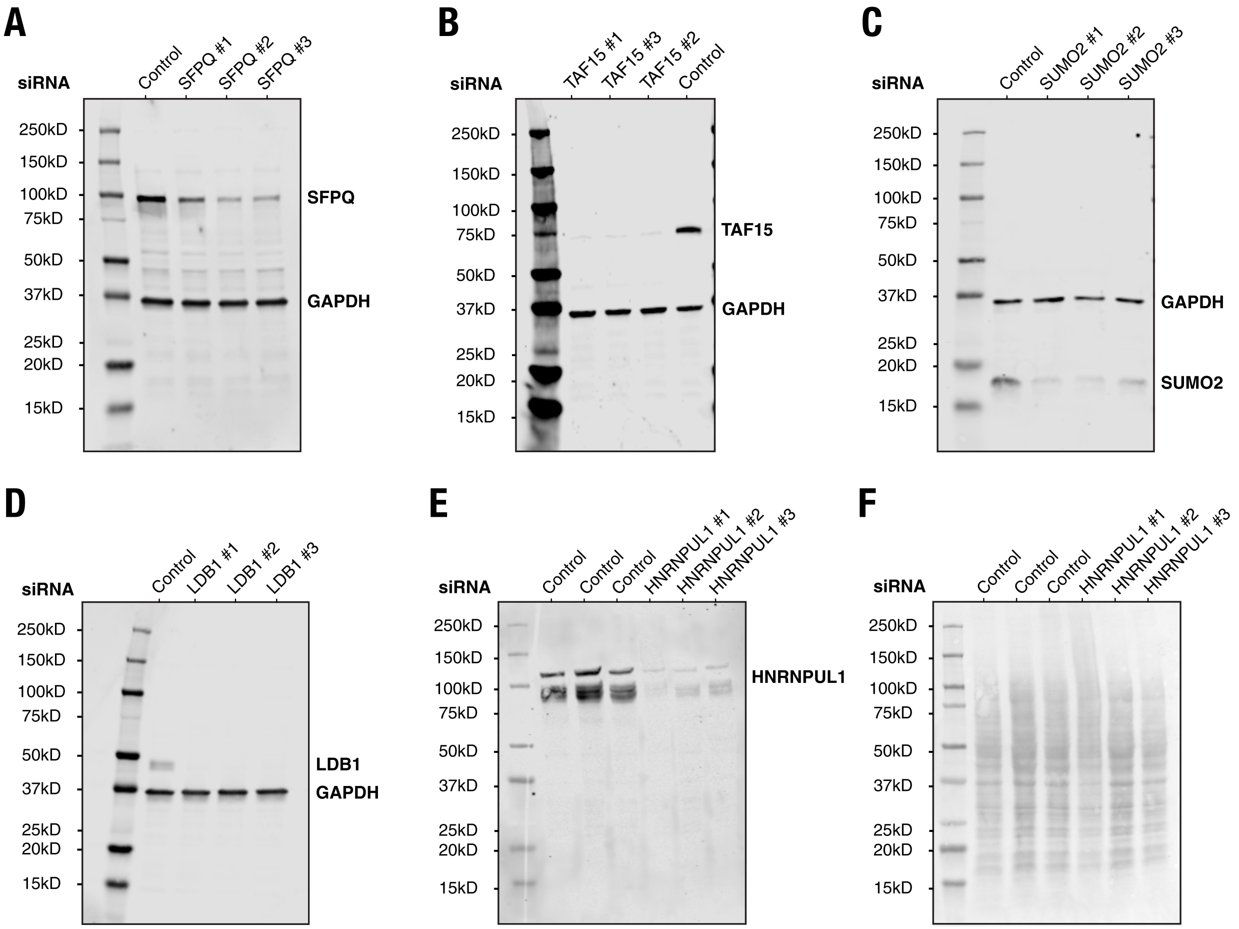


**Fig. S7. Immunoblot analyses that confirm siRNA-mediated knockdown of PhaseID-identified partners in A673 cells.**

(A-D) Immunoblots of SFPQ (A), TAF15 (B), SUMO2 (C), and LDB1 (D) from whole-cell lysates of A673 cells transfected with respective siRNAs or negative control siRNAs with no specific target in the cell. GAPDH served as a loading control. For each gene, siRNA numbers (#1, #2, #3) indicate distinct siRNA oligos targeting different regions of the same gene.

(E-F) Immunoblots of HNRNPUL1 (E) from whole-cell lysates of A673 cells transfected with HNRNPUL1 siRNAs or negative control siRNAs. All lanes were loaded with equal amounts of total protein, as shown by Ponceau S staining of the same membrane (F). The knockdown efficiency for SFPQ, TAF15, SUMO2, LDB1, and HNRNPUL1 was assessed 72 hrs after siRNA transfection.

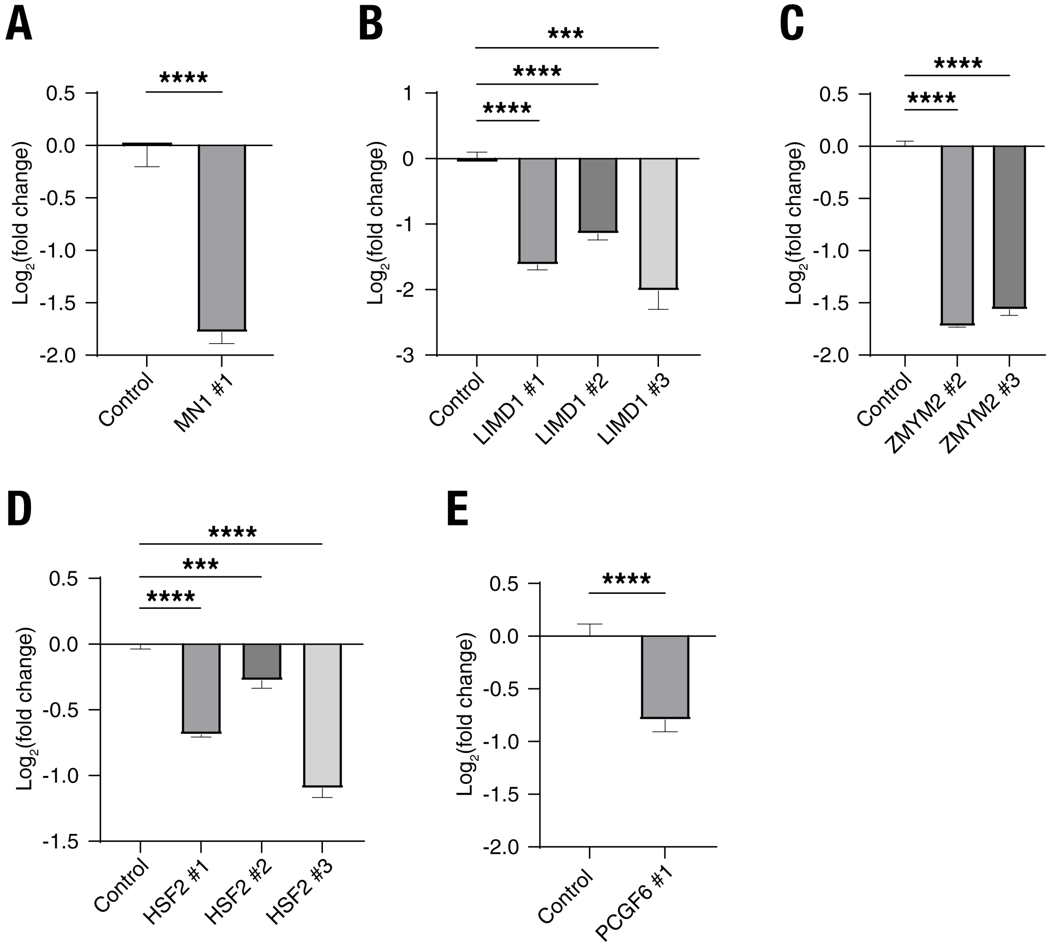


**Fig. S8. RT-qPCR analyses that confirm siRNA-mediated knockdown of PhaseID-identified partners in A673 cells.**

The evaluated knockdowns are for MN1 (A), LIMD1 (B), ZMYM2 (C), HSF2 (D), and PCGF6 (E). Cells transfected with a siRNA with no specific target in the cell were used as the control. For each gene, siRNA numbers (#1, #2, #3) indicate distinct siRNA oligos targeting different regions of the same gene. Error bars represent standard deviations of four technical replicates. Statistical significance: $p < 0.001$ (***), $p < 0.0001$ (****); Welch's two-sample $t$-test.

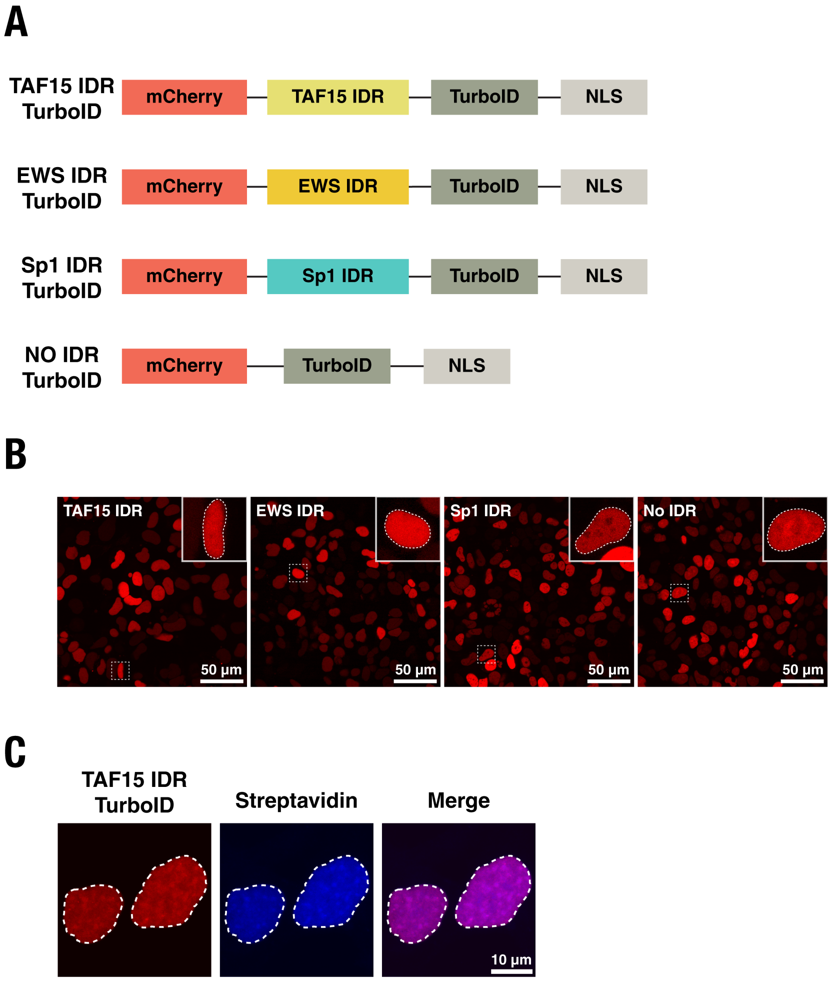


**Fig. S9. Construct design and characterization data for conventional TurboID-mediated proximity labeling.**
(A) Schematics of the TurboID constructs for TAF15, EWS, and Sp1 IDRs, as well as the no IDR control.
(B) Confocal fluorescence images of modified HEK293 cells expressing TurboID constructs for different IDRs and the no IDR control. Scale bars, 50 µm.
(C) Confocal fluorescence images of modified HEK293 cells expressing the mCherry-labeled TAF15 IDR TurboID construct (red) after the cells were treated with biotin (50 µM, 10 min) and stained with AF750-labeled streptavidin (blue). The biotinylated proteins are distributed nearly homogeneously with no detectable puncta throughout the nuclei. A white dashed contour outlines the cell nucleus. Scale bar, 10 µm.

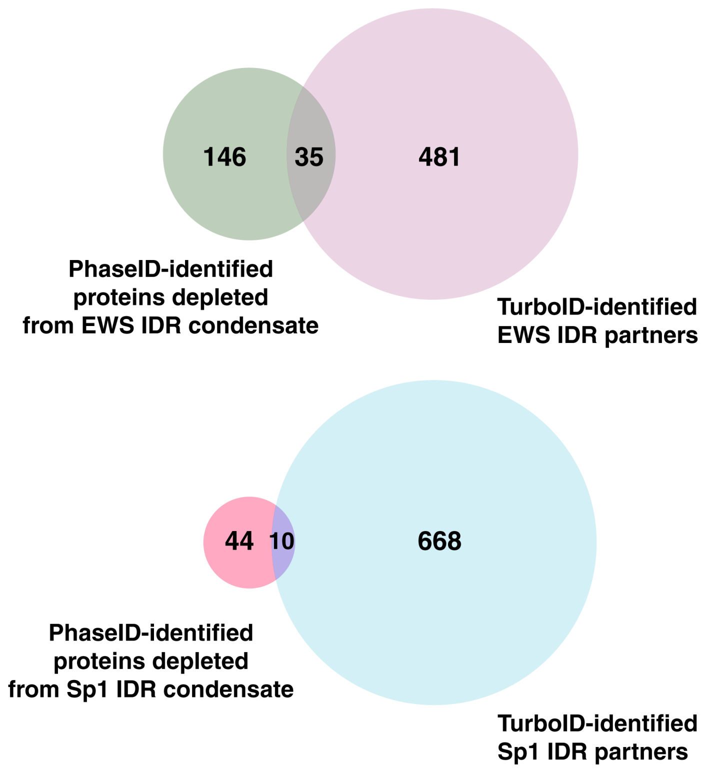


**Fig. S10. Venn diagrams comparing PhaseID-identified proteins depleted from the condensates of an IDR and TurboID-identified partners of the IDR.**
For both EWS IDR (top) and Sp1 IDR (bottom), a subset of TurboID-detected IDR partners are excluded from the induced condensates of the target IDR as determined by PhaseID.

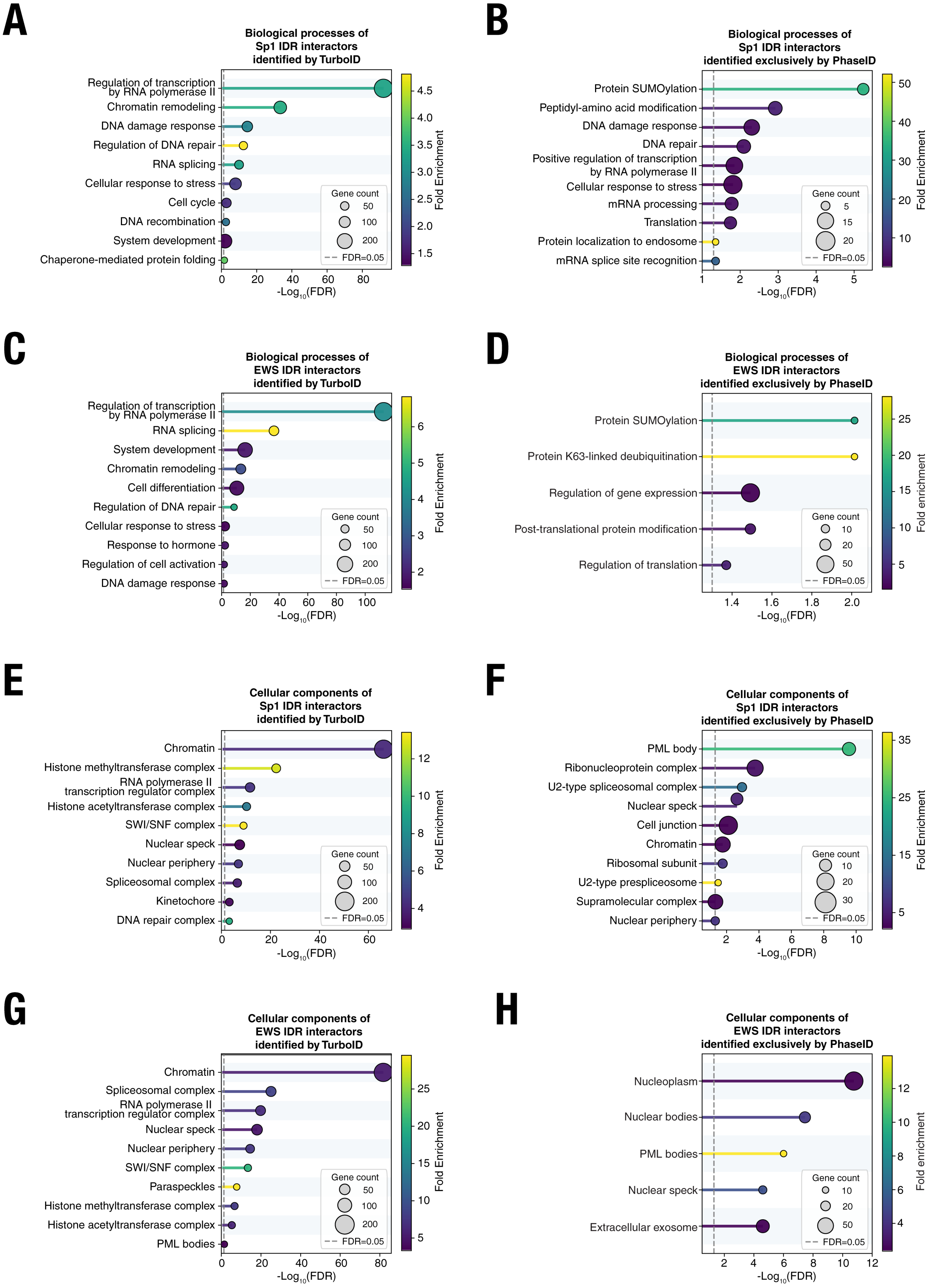
A
Biological processes of
Sp1 IDR interactors
identified by TurboID
Regulation of transcription
by RNA polymerase II
Chromatin remodeling
DNA damage response
Regulation of DNA repair
RNA splicing
Cellular response to stress
Cell cycle
DNA recombination
System development
Chaperone-mediated protein folding
Gene count
50
100
200
FDR=0.05
-Log10(FDR)
Fold Enrichment
B
Biological processes of
Sp1 IDR interactors
identified exclusively by PhaseID
Protein SUMOylation
Peptidyl-amino acid modification
DNA damage response
DNA repair
Positive regulation of transcription
by RNA polymerase II
Cellular response to stress
mRNA processing
Translation
Protein localization to endosome
mRNA splice site recognition
Gene count
5
15
20
FDR=0.05
-Log10(FDR)
Fold Enrichment
C
Biological processes of
EWS IDR interactors
identified by TurboID
Regulation of transcription
by RNA polymerase II
RNA splicing
System development
Chromatin remodeling
Cell differentiation
Regulation of DNA repair
Cellular response to stress
Response to hormone
Regulation of cell activation
DNA damage response
Gene count
50
100
200
FDR=0.05
-Log10(FDR)
Fold Enrichment
D
Biological processes of
EWS IDR interactors
identified exclusively by PhaseID
Protein SUMOylation
Protein K63-linked deubiquitination
Regulation of gene expression
Post-translational protein modification
Regulation of translation
Gene count
10
20
50
FDR=0.05
-Log10(FDR)
Fold enrichment
E
Cellular components of
Sp1 IDR interactors
identified by TurboID
Chromatin
Histone methyltransferase complex
RNA polymerase II
transcription regulator complex
Histone acetyltransferase complex
SWI/SNF complex
Nuclear speck
Nuclear periphery
Spliceosomal complex
Kinetochore
DNA repair complex
Gene count
50
100
200
FDR=0.05
-Log10(FDR)
Fold Enrichment
F
Cellular components of
Sp1 IDR interactors
identified exclusively by PhaseID
PML body
Ribonucleoprotein complex
U2-type spliceosomal complex
Nuclear speck
Cell junction
Chromatin
Ribosomal subunit
U2-type prespliceosome
Supramolecular complex
Nuclear periphery
Gene count
10
20
30
FDR=0.05
-Log10(FDR)
Fold Enrichment
G
Cellular components of
EWS IDR interactors
identified by TurboID
Chromatin
Spliceosomal complex
RNA polymerase II
transcription regulator complex
Nuclear speck
Nuclear periphery
SWI/SNF complex
Paraspeckles
Histone methyltransferase complex
Histone acetyltransferase complex
PML bodies
Gene count
50
100
200
FDR=0.05
-Log10(FDR)
Fold Enrichment
H
Cellular components of
EWS IDR interactors
identified exclusively by PhaseID
Nucleoplasm
Nuclear bodies
PML bodies
Nuclear speck
Extracellular exosome
Gene count
10
20
50
FDR=0.05
-Log10(FDR)
Fold enrichment

**Fig. S11. Comparison of Gene Ontology enrichment analyses of TurboID- and PhaseID-identified interactomes of Sp1 and EWS IDRs.**

(A–D) Biological Process terms enriched among the TurboID-identified partners of Sp1 IDR (A) and EWS IDR (C) or Biological Process terms enriched among the partners of Sp1 IDR (B) and EWS IDR (D) identified exclusively by PhaseID.

(E–H) Cellular Component terms enriched among the TurboID-identified partners of Sp1 IDR (E) and EWS IDR (G) or Biological Process terms enriched among the partners of Sp1 IDR (F) and EWS IDR (H) identified exclusively by PhaseID.

The top significantly enriched GO terms are ranked based on the values of −log10(FDR), with the circle size indicating the gene count and the color scale representing the fold enrichment. The dashed vertical line denotes the cutoff for significant enrichment (FDR<0.05). FDR (false discovery rate) represents the multiple-testing-adjusted p-value, as calculated by the DAVID Bioinformatics Resources (*63*).

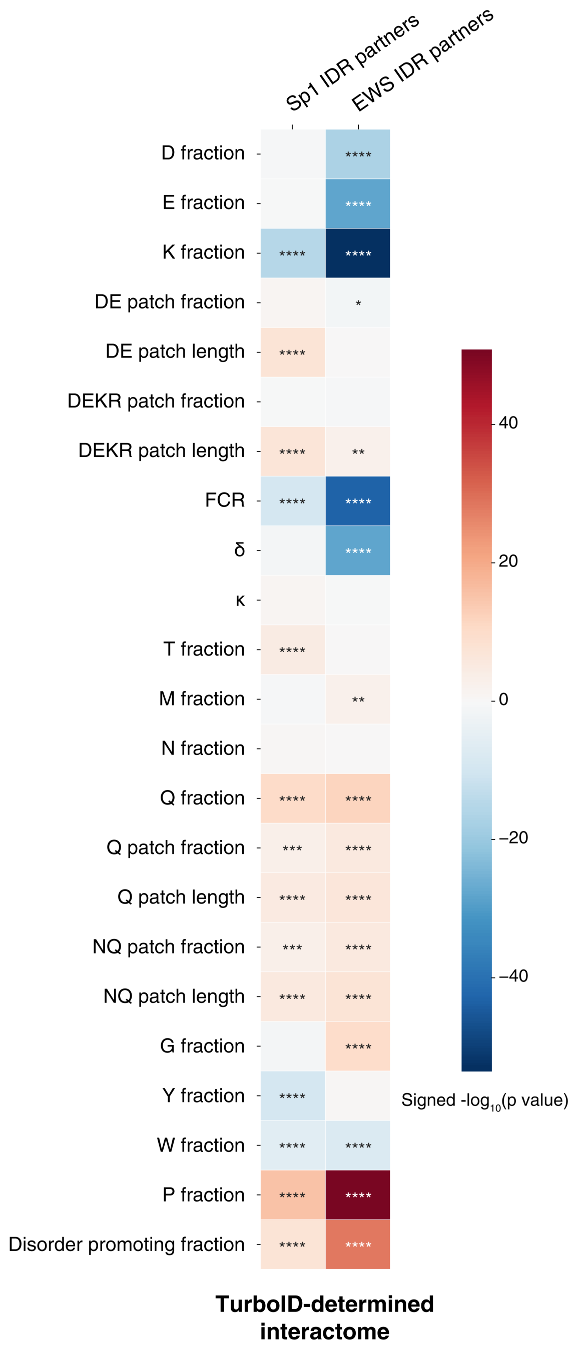


**Fig. S12. Sequence feature analysis of the TurboID-determined interactomes of EWS and Sp1 IDRs.**

Heatmap of signed *p* values showing that TurboID-identified protein partners of EWS IDR and Sp1 IDR have specific sequence parameter values distinct from those of the mass spectrometry-detected proteome. For each sequence parameter, values for individual TurboID-identified proteins were normalized to z-scores using the proteome mean and standard deviation. Each z-score represents how far a protein's parameter value deviates from the proteome average. In the plot, a positive value (red) or a negative value (blue) denote that the sequence parameter has a higher or lower mean z-score for the protein group, respectively, compared to the proteome. The sign reflects the direction of the mean z-score difference, and the magnitude corresponds to $-\log_{10}$(p-value). The sequence parameter values were computed with localCIDER (*94*). FCR, fraction of charged residues. κ quantifies the linear patterning of charged residues along a protein sequence. δ is the variance of local charge asymmetry from the overall charge asymmetry across defined residue windows in a protein sequence. Patches are contiguous runs of ≥5 identical or chemically similar residues. See supplemental methods for details. Statistical significance: $p < 0.05$ (*), $p < 0.01$ (**), $p < 0.001$ (***), $p < 0.0001$ (****); Welch's two-sample *t*-test.

**Table S1. Antibodies and Fluorescent Probes**

| Antibodies | Source | Concentration | Host |
|---|---|---|---|
| Streptavidin, Alexa Fluor 750 conjugate | Thermo Fisher, S21384 | 1:50 (IF) | - |
| IRDye 800CW Streptavidin | Licor Bio, 926-32230 | 1:3000 (WB) | - |
| anti-GAPDH | Santa Cruz Biotechnology, sc-47724 | 1:1000 (WB) | Mouse |
| anti-GAPDH | Cell signaling technology, 2118S | 1:1000 (WB) | Rabbit |
| anti-mCherry | Abcam, ab167453 | 1:1000 (WB) | Rabbit |
| anti-GFP | Thermo Fisher, A-11122 | 1:2000 (WB) | Rabbit |
| anti-YAP1 | Santa Cruz Biotechnology, sc-101199 | 1:50 (IF) | Mouse |
| anti-EWSR1 | Abcam, ab54708 | 1:200 (IF) | Mouse |
| anti-SFPQ | Santa Cruz Biotechnology, sc-101137 | 1:50 (IF)/1:1000 (WB) | Mouse |
| anti-SUMO2 | Thermo Fisher, MA5-37627 | 1:25 (IF)/1:1000 (WB) | Mouse |
| anti-BRCA1 | Thermo Fisher, MA1-23164 | 1:100 (IF) | Mouse |
| anti-CDCA2 | Thermo Fisher, PA5-55299 | 1:25 (IF) | Rabbit |
| anti-TAF15 | Abcam, ab134916 | 1:10000 (WB) | Rabbit |
| anti-LDB1 | Thermo Fisher, PA5-56948 | 1:1000 (WB) | Rabbit |
| anti-HNRNPUL1 | Santa Cruz Biotechnology, sc-393975 | 1:1000 (WB) | Mouse |
| F(ab')2-Rabbit anti-Mouse IgG (H+L) Cross-Adsorbed Secondary Antibody, Alexa Fluor 488 | Thermo Fisher, A-21204 | 1:1000 (IF) | Rabbit |
| Donkey anti-Rabbit IgG (H+L) Highly Cross-Adsorbed Secondary Antibody, Alexa Fluor 488 | Thermo Fisher, A-21206 | 1:1000 (IF) | Donkey |
| IRDye 680RD Goat anti-Rabbit IgG Secondary Antibody | Licor Bio, 926-68071 | 1:15000 (WB) | Goat |
| IRDye 800CW Goat anti-Mouse IgG Secondary Antibody | Licor Bio, 926-32210 | 1:15000 (WB) | Goat |

**Table S2. Silencer Select siRNA IDs from Thermo Fisher**

| Name | Gene ID | siRNA ID |
|---|---|---|
| Negative control No.2 | | s814 |
| HNRNPUL1 #2 | 11100 | s21884 |
| HNRNPUL1 #3 | 11100 | s21885 |
| SFPQ #1 | 6421 | s12710 |
| SFPQ #2 | 6421 | s12711 |
| SFPQ #3 | 6421 | s12712 |
| TAF15 #3 | 8148 | s15658 |
| LDB1 #1 | 8861 | s16921 |
| LDB1 #2 | 8861 | s16922 |
| LDB1 #3 | 8861 | s16923 |
| SUMO2 #2 | 6613 | s13180 |
| SUMO2 #3 | 6613 | s13181 |
| MN1 #1 | 4330 | s8895 |
| LIMD1 #1 | 8994 | s17158 |
| LIMD1 #2 | 8994 | s17159 |
| LIMD1 #3 | 8994 | s17160 |
| ZMYM2 #2 | 7750 | s15268 |
| ZMYM2 #3 | 7750 | s15266 |
| HSF2 #1 | 3298 | s6954 |
| HSF2 #2 | 3298 | s6955 |
| HSF2 #3 | 3298 | s6953 |
| PCGF6 #1 | 84108 | s38515 |

**Table S3. Sequences of primers used in RT-qPCR**

| Name | Sequence (5’-3’) |
|---|---|
| Forward primer targeting *CYP4F22* | ATATCCGAGCCGAAGCAGACAC |
| Reverse primer targeting *CYP4F22* | CGGCATTTCTCCTGGTATTCCG |
| Forward primer targeting *CCK* | TGAGGGTATCGCAGAGAACGGA |
| Reverse primer targeting *CCK* | CGGTCACTTATCCTGTGGCTGG |
| Forward primer targeting *PPP1R1A* | CACAGAAGTGGAGTCAAGGCTG |
| Reverse primer targeting *PPP1R1A* | TTGGCTCCCTTGGAATCCAGTG |
| Forward primer targeting *NR0B1* | AGGGGACCGTGCTCTTTAAC |
| Reverse primer targeting *NR0B1* | CTGAGTTCCCCACTGGAGTC |
| Forward primer targeting *BIRC5* | TGCCCCGACGTTGCC |
| Reverse primer targeting *BIRC5* | CAGTTCTTGAATGTAGAGATGCGGT |
| Forward primer targeting *B4GALT5* | TCCTCGCTGCTGTACTTCG |
| Reverse primer targeting *B4GALT5* | AATGCCTTGGGCTTGCATCA |
| Forward primer targeting *MMP14* | GCAGAAGTTTTACGGCTTGCAA |
| Reverse primer targeting *MMP14* | CCTTCGAACATTGGCCTTGAT |
| Forward primer targeting *IGF1R* | GAAGTCTGGCTCCGGAGGAGGGTC |
| Reverse primer targeting *IGF1R* | ATGTGGAGGTAGCCCTCGATCAC |
| Forward primer targeting *GAPDH* | GGTCTCCTCTGACTTCAACA |
| Reverse primer targeting *GAPDH* | GTGAGGGTCTCTCTCTTCCT |
| Forward primer targeting *MN1* | CAGAACCCCAACAGCAAAGAA |
| Reverse primer targeting *MN1* | GACAGACAGGCACTGCAAGTG |
| Forward primer targeting *LIMD1* | TGGGGAACCTCTACCATGAC |
| Reverse primer targeting *LIMD1* | CACAAAACACTTTGCCGTTG |
| Forward primer targeting *ZMYM2* | GGTAAACTAACTGAGATTCGCCA |
| Reverse primer targeting *ZMYM2* | CCAGAACATTATTTCCAGCACCT |
| Forward primer targeting *HSF2* | ATTCAGAGTGGAGAGCAGAATG |
| Reverse primer targeting *HSF2* | GACAGCACTAGACATGAGA |
| Forward primer targeting *PCGF6* | TGTTCCACAGCCAGTCCCTT |
| Reverse primer targeting *PCGF6* | TGCACGTCGGATTTCCCTTA |
